# Dissociation Protocols used for Sarcoma Tissues Bias the Transcriptome observed in Single-cell and Single-nucleus RNA sequencing

**DOI:** 10.1101/2022.01.21.476982

**Authors:** Danh D. Truong, Salah-Eddine Lamhamedi-Cherradi, Robert W. Porter, Sandhya Krishnan, Jyothishmathi Swaminathan, Amber Gibson, Alexander J. Lazar, John A. Livingston, Vidya Gopalakrishnan, Nancy Gordon, Najat C. Daw, Richard Gorlick, Joseph A. Ludwig

**Author notes:** **Corresponding author** Correspondence to Joseph A. Ludwig.

## Abstract

**Background:** Single-cell RNA-seq has emerged as an innovative technology used to study complex tissues and characterize cell types, states, and lineages at a single-cell level. Classification of bulk tumors by their individual cellular constituents has also created new opportunities to generate single-cell atlases for many organs, cancers, and developmental models. Despite the tremendous promise of this technology, recent evidence studying epithelial tissues and diverse carcinomas suggests the methods used for tissue processing, cell disaggregation, and preservation can significantly bias gene expression and alter the observed cell types. To determine whether sarcomas – tumors of mesenchymal origin – are subject to the same technical artifacts, we profiled patient-derived tumor explants (PDXs) propagated from three aggressive subtypes: osteosarcoma, Ewing sarcoma (ES), desmoplastic small round cell tumor (DSRCT). Given the rarity of these sarcoma subtypes, we explored whether single-nuclei RNA-seq from more widely available archival frozen specimens could accurately be identified by gene expression signatures linked to tissue phenotype or pathognomonic fusion proteins.

**Results:** We systematically assessed dissociation methods across different sarcoma subtypes. We compared gene expression from single-cell and single-nucleus RNA-sequencing of 125,831 whole-cells and nuclei from ES, DSRCT, and osteosarcoma PDXs. We detected warm dissociation artifacts in single-cell samples and gene length bias in single-nucleus samples. Classic sarcoma gene signatures were observed regardless of dissociation method. In addition, we showed that dissociation method biases can be computationally corrected.

**Conclusions:** We highlighted transcriptional biases, including warm dissociation and gene-length biases, introduced by the dissociation method for various sarcoma subtypes. This work is the first to characterize how the dissociation methods used for sc/snRNA-seq may affect the interpretation of the molecular features in sarcoma PDXs.

## Background

Tumors are composed of a diverse multicellular microenvironment that dictate cancer progression and response to therapy. While cells share an identical genome, their phenotype and behavior are driven by their transcriptome and proteome^1^. Cellular heterogeneity within the tumor ecosystem has precluded the ability to fully understand the cell biology and interactions that drive cancer progression^1^. Recently, single-cell RNA-seq (scRNA-seq) has emerged as an innovative technology to characterize individual cells from heterogeneous tissues in order to understand cell types, states, and lineages^2^. Rapid adoption of this technology has led to a flurry of research generating single-cell atlases for many organs, cancers, and developmental models enriching our understanding of cell biology^3^.

Despite the tremendous success of this technology when applied to different cancer types, sarcomas, which are cancers of mesenchymal origin, have not yet widely benefited from the adoption of scRNA-seq. Differences in tissue origin may require optimized dissociation to capture accurate *in vivo* gene expression and cellular composition. Further, the enzymatic and mechanical methods used to dissociate cells are known to bias cellular composition and reduce cellular quality. Many gold standard dissociation protocols require extended incubation at 37°C, where cellular transcription is still active and may introduce gene expression artifacts^4^. Cold-active protease is a recent alternative to dissociation at 37°C, which may limit and minimize transcriptional activity and environmental stresses on cells^4,5^.

Challenges in obtaining fresh clinical specimens and the logistical issues to immediately process specimens have also hindered workflows^6^. While cancer models for sarcoma, including cell lines, xenografts, and PDXs, are readily accessible for scRNA-seq, the extent that these models represent the original cancer specimen have not yet been adequately evaluated. Single-nucleus RNA-seq (snRNA-seq) of accessible frozen tissue has demonstrated concordance with scRNA-seq^6-10^. SnRNA-seq can remove the limitations for obtaining fresh tissue and immediate processing by enabling access to archival tissue and ease the coordination of tissue acquisition by allowing sequencing of snap-frozen tissue. Furthermore, difficulties with cell fragility or size when considering scRNA-seq can be circumvented using snRNA-seq.

The biases introduced by different methods have been studied between single-cell and single-nucleus as well as dissociation using cold-active proteases and standard digestion at 37°C^4^. However, these studies did not include sarcoma specimens, which differ significantly from epithelial tissues and carcinomas in their expression not only by lineage but also integrins and cell-cell adhesions^11,12^. To fully realize the potential of scRNA-seq and snRNA-seq in three of the fifty or more unique sarcoma subtypes, we systematically assessed the effect temperature has upon enzymatic dissociation of fresh tissue and, secondarily, studied whether snRNA-seq maintains key transcriptomic profiles determined using scRNA-seq. We focused our analysis on well-controlled PDX specimens of different and rare sarcomas to enable sample accessibility since fresh sarcoma specimens are difficult to acquire. This further enabled our group to explore multiple dissociation methods on the same sample.

Though more than fifty distinct sarcoma subtypes exist, our work takes an important step to layout the technical and analytical framework needed for scRNA-seq and snRNA-seq analysis of osteosarcoma, ES, and DSRCT, three highly aggressive sarcoma samples that affect adolescents and young adults. Our work highlights notable method-dependent biases, as well as computational tools used to remove them when rare archival frozen samples are assessed by snRNA-seq.

## Results

### Single-cell and single-nucleus RNA sequencing of sarcoma subtypes

In this work, we studied sarcomas from varying tissue origins, including osteosarcoma (OS), Ewing’s sarcoma (ES), and desmoplastic small round cell tumor (DSRCT) (**Fig. 1**). We used different dissociation protocols: Miltenyi Tumor Dissociation Kit, cold-active protease derived from *Bacillus licheniformis*, and Nuclei EZ Prep. These three protocols are described herein as Warm, Cold, and Nuclei protocols. For the OS specimens, we used the same Nuclei protocol and a different Warm protocol optimized for OS.

**Figure 1.**
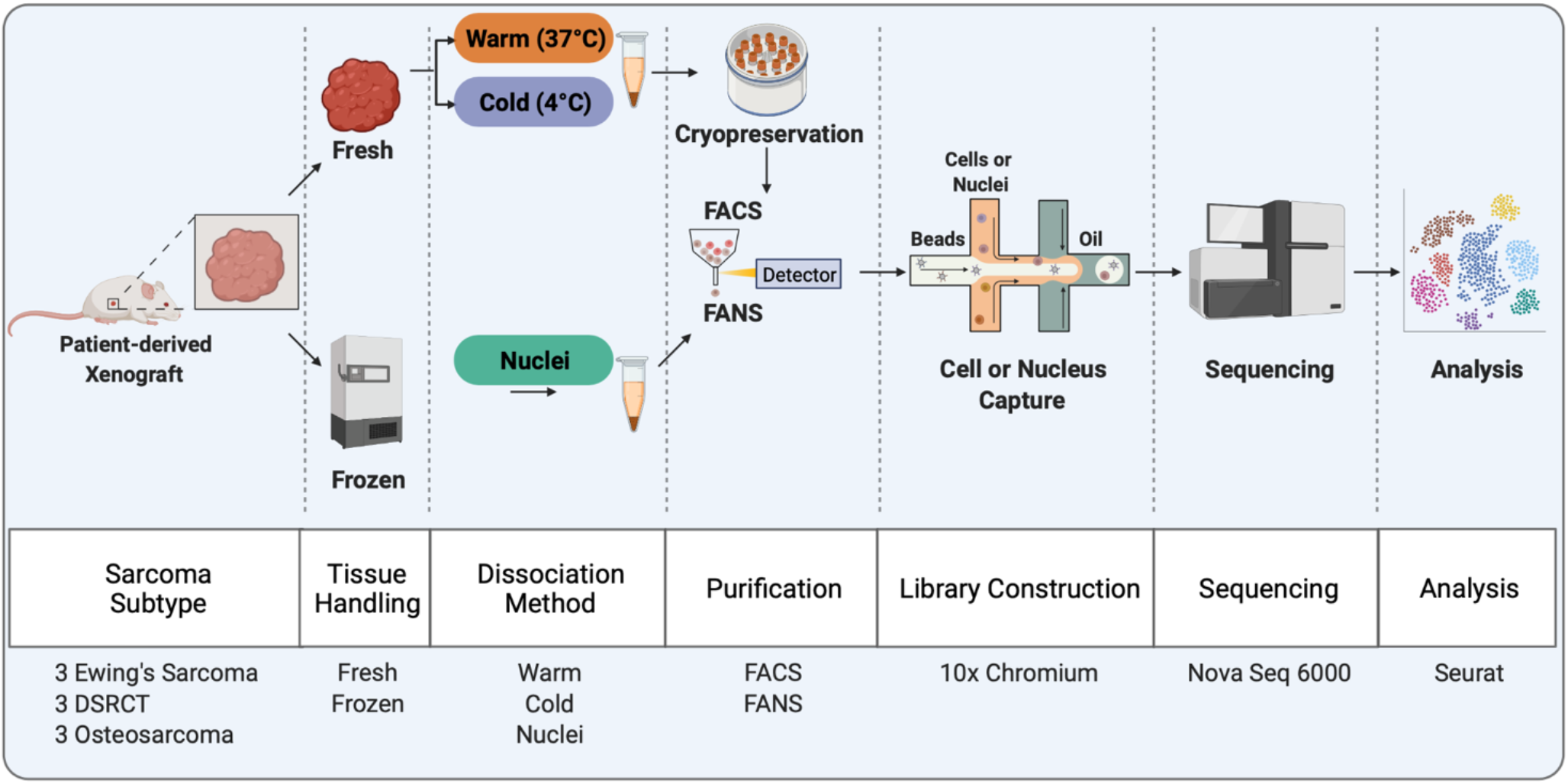
Overview of Workflow and Experiments in this study. Schematic depicting the sarcoma subtypes used in this study and the general workflow for the specific experiments. FACs: fluorescence-activated cell sorting, FANS: fluorescence-activated nucleus sorting, DSRCT: desmoplastic small round cell tumor

For DSRCT and ES specimens, we performed the additional Cold protocol, using the cold-active protease, as we had more specimens available. Each sarcoma subtype included three PDX specimens derived from different patients. In total, we analyzed 125,831 whole-cells and nuclei across the three sarcoma subtypes and three dissociation protocols.

### Evaluation of quality control metrics for tissue dissociation protocols

Previous work has shown that dying and dead cells can influence the transcriptome and introduce artifacts that preclude useful biological insight^4^. To evaluate this effect in sarcoma, we evaluated and compared several protocols based on cell/nucleus quality and transcriptomic signatures. For cell/nucleus quality, we measured the percent of reads mapping to the transcriptome, number of genes, unique molecular identifiers (UMIs), and percent of mitochondrial genes for each cell or nucleus. We optimized our strategy to enrich for live single cells and single nucleus, respectively, by incorporating fluorescently activated cell sorting (FACS) or fluorescently activated nuclei sorting (FANS) prior to sequencing (**Fig. S1**).

Next, we evaluated common quality control (QC) metrics across all samples to assess the effect of each dissociation protocol (**Fig. 2A**). We observed some variations in QC metrics for the number of genes and UMIs when comparing between protocols while limiting the comparisons to between PDXs of each sarcoma subtype. However, some of the variations could be explained by number of cells sequenced and sequencing depth since there is an inverse relationship between these two metrics when total reads are kept constant (**Fig. 2B**). Expectedly, nucleus samples demonstrated little to no percentage of mitochondrial genes since purified nuclei do not contain mitochondrial transcripts. With respect to each protocol, we did not discern a positive or negative influence on the QC metrics.

**Figure 2.**
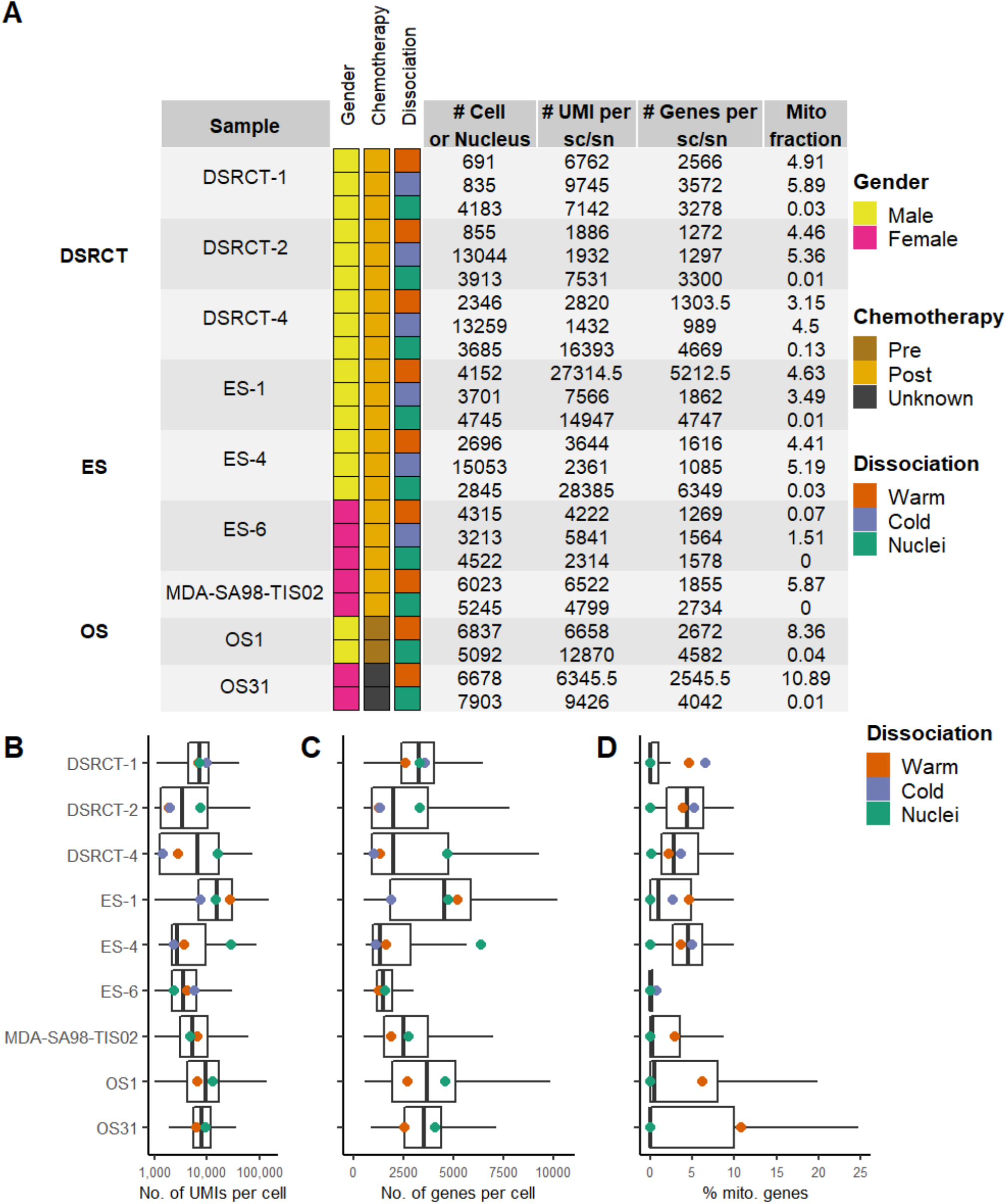
Patient information and quality control metrics. **a** Overview of all sarcoma subtypes that were processed and evaluated. For each sample, the number of cells or nuclei passing QC thresholds, number of sequencing reads per cell/nuclei, number of genes per cell/nuclei, and the median percentage of UMIs mapping to mitochondrial genes are displayed in the table. All samples had less than 0.01 doublet fraction. **b** Distributions (median and first and third quartiles) for number of UMIs per cells/nuclei, number of genes per cell/nuclei, and percentage of UMIs mapping to mitochondrial genes vary across sarcoma subtypes and choice of protocol.

### Dissociation protocol biases the transcriptome

To determine whether protocol-specific differences in gene expression exist, we visualized the UMAP embeddings of all whole-cells and nuclei without batch or technical corrections. When colored by sarcoma subtype, the same sarcoma subtype cluster together but with two distinct clusters for each sarcoma subtype except for OS (**Fig. 3A**). We suspect that this may be due to biases from the different dissociation protocols. When labeled by fresh specimens (whole cells) or frozen specimens (nuclei), we identified a distinct delineation between fresh and frozen tissues in the UMAP. The observed differences within the UMAP, we hypothesized, stem from biological artifacts linked to fresh tissue dissociation or technical artifacts that reflect a core set of mRNA transcripts preferentially retained within the nucleus. By coloring the UMAP embedding by dissociation type, cells processed using the Warm and Cold methods partially overlapped for each PDX, whereas Nuclei clusters remained segregated.

**Figure 3.**
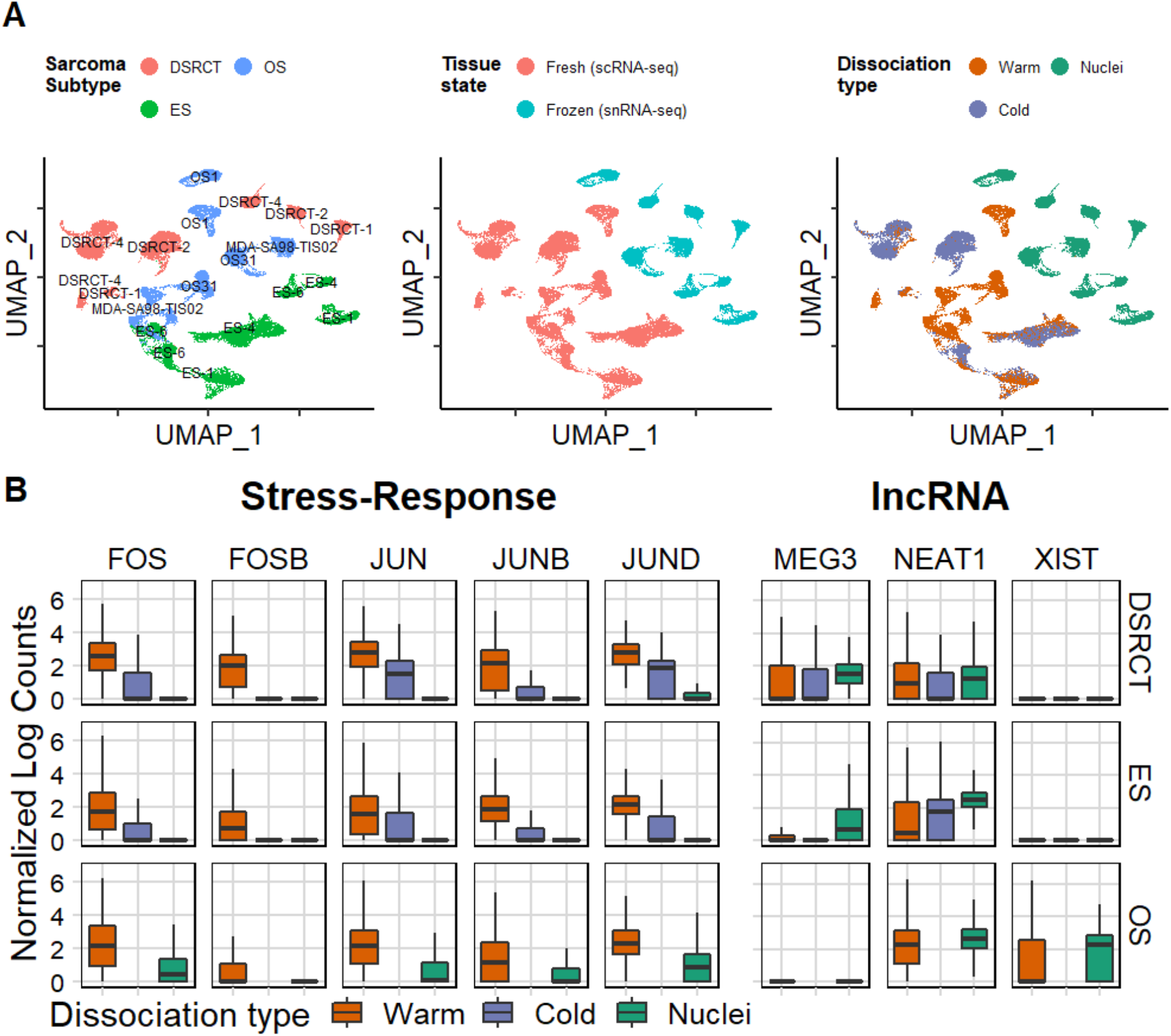
Performance of each protocol on various sarcoma subtypes. **a** UMAP embedding of all cells labeled by Sarcoma subtype, Tissue state, and Dissociation type. **b** Enrichment of select stress-response genes in the Warm dissociation protocol and select lncRNA in the Nuclei isolation protocol.

Previous reports using normal epithelial tissues and carcinomas revealed that warm enzymatic dissociation (i.e., at 37°C) invoked a distinct ‘Warm Dissociation Signature’ enriched in FOS, FOSB, and JUN^7^. To investigate if similar dissociation-specific biases occur in sarcomas exposed to collagenase at 37°C, we selected a partial list of the top genes within the Warm Dissociation signature and compared their expression. Averaged gene expression from each sarcoma subtype showed that these genes are, indeed, elevated in the Warm protocol (**Fig. 3B**). Furthermore, since prior literature have stated that long non-coding RNAs (lncRNAs) are localized to the nucleus, we also explored their gene expression in these specimens^13^. Consistent with previous results, we observed that lncRNAs were elevated in the Nuclei protocol. Together, these results indicate the method chosen dissociation has a profound effect on gene expression.

To further characterize how scRNA-seq and snRNA-seq affect transcript abundance, we performed an analysis of differentially expressed genes (DEGs). Warm and Nuclei protocols demonstrated a consistent trend for each sarcoma type. Genes with the largest fold-change in the Warm method included mitochondrial and ribosomal protein genes (**Fig. 4, Table S1-3**). This was expected since the mitochondria (and their innate transcripts) are removed entirely during the Nuclei dissociation method. Similarly, enrichment of ribosomal protein genes was also noted in a comparison between scRNA-seq and snRNA-seq for kidney tissue^14^. On the other hand, genes enriched in the Nuclei protocol did not have a clear consensus or overlap between sarcoma types. Analyzing the DEGs in the Warm protocol, we found a common set of 325 genes enriched after filtering for log2 fold change over 1.5. Similarly, we found 117 genes enriched in the Nuclei protocol (**Fig. S2A, B**). Next, we performed a pathway enrichment of the MSigDB hallmark gene set using Enrichr. We observed stress-associated pathways in each sarcoma type that was enriched in the Warm protocol including Hypoxia, Apoptosis, DNA repair, and TNF-alpha Signaling via NF-kB, which is consistent with prior work^4^. On the other hand, for the Nuclei protocol, we observed enrichment in Mitotic Spindle. When comparing the Warm and Cold protocols for only ES and DSRCT, we again observed an increase in several of the commonly identified stress-related pathways like previous results (**Fig. S3, Table S4, 5**). The UMAP embedding suggested that the differences in Warm and Cold are minimal due to the two data sets overlapping when accounting for each PDX (**Fig. 3A**). Furthermore, we found in total 24 commonly enriched pathways suggesting a core set of conserved genes enriched in the Warm protocol for sarcoma samples (**Fig. S2C**). Interestingly, we did not observe any common pathways enriched between sarcomas for the Nuclei protocol.

**Figure 4.**
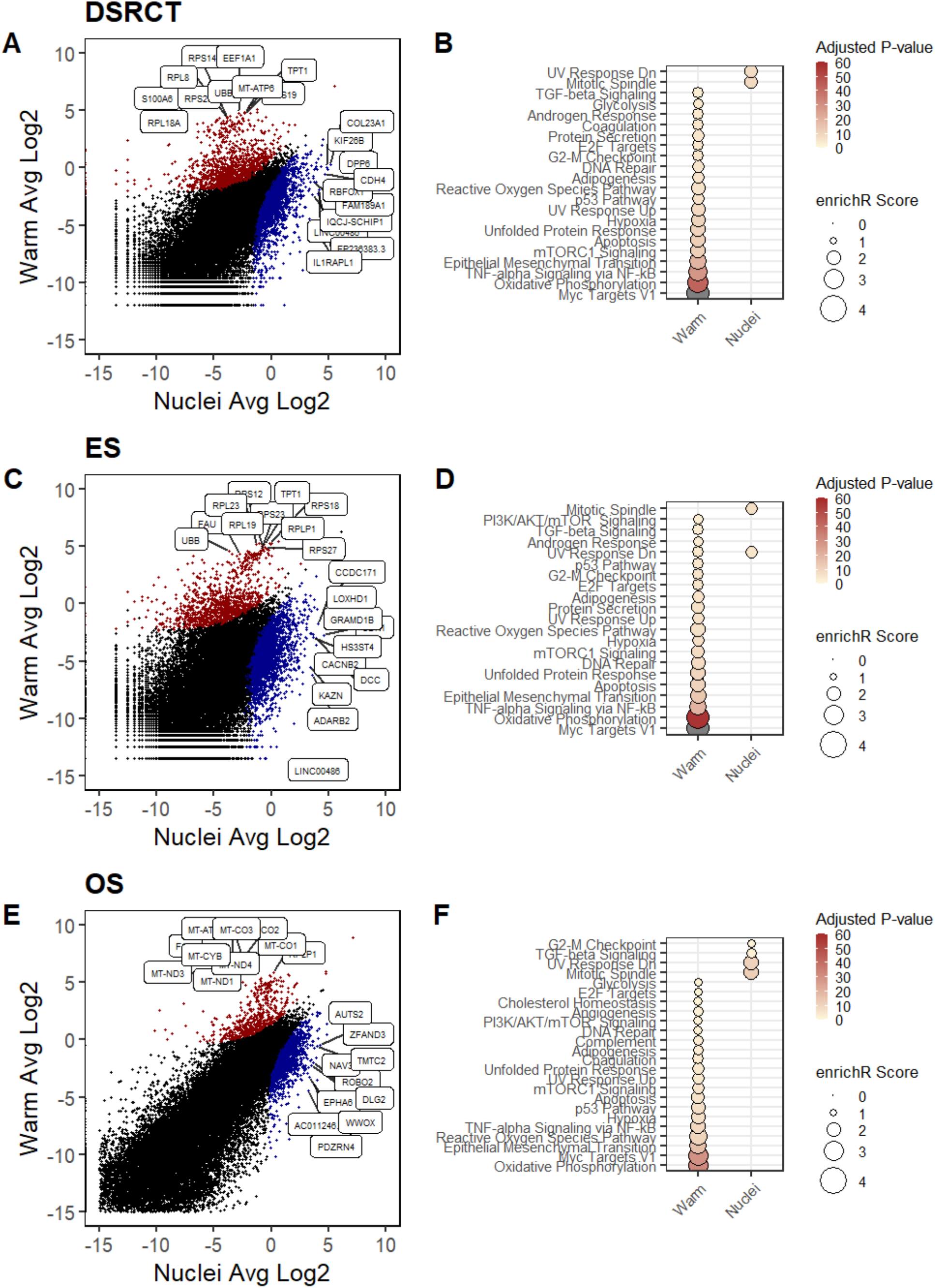
DEG Biases introduced by Warm and Nuclei Protocols. Scatter plot of log transformed gene expression levels between warm and nuclei on the left column. Red indicates up-regulated in warm, and blue indicates up-regulated in nuclei with p-value < 0.05. Black is non-significant. Dot plot of enrichR scores of the Hallmark gene sets from MSigDB are on the right column. Plots are shown for DSRCT **a, b**; ES **c, d**; and OS **e, f**.

### Sarcoma signatures are preserved irrespective of the method used for dissociation

Next, to evaluate if any of the protocols influenced signatures associated with a particular sarcoma type, we analyzed expression of gene sets curated from literature (**Table S6**). For ES, we used a set of genes that are direct targets of the EWS-FLI1 fusion protein, which included KDSR, CAV1 and FCGRT^15^. Likewise, a gene set for EWS-WT1 targets, generated from cell lines, was used to evaluate the effect of each protocol in DSRCT^16^. Since OS lacks a clearly defined gene set, and often contains cells of partial fibroblastic, chondroblastic, or osteoblastic lineage commitment, we utilized curated genes associated with osteoblastic and chondroblastic signatures classically associated with the putative tissue origin of OS. Strikingly, the unique sarcoma subtype-specific gene signatures were preserved across all dissociation protocols. (**Fig. 5**). This suggests that regardless of dissociation protocol biases, the cells still exhibit the classic signatures for each sarcoma studied. For instance, the EWS-WT1 gene targets are upregulated in only the DSRCT PDX specimens. Likewise, the EWS-FLI1 target genes are only enriched in ES, irrespective of protocol used. However, when comparing between protocols for ES, we observed overexpression of the EWS-FLI1 gene set in the Nuclei protocol. While we did not observe this phenomenon in the other gene sets, we explored the idea of a Nuclei protocol bias.

**Figure 5.**
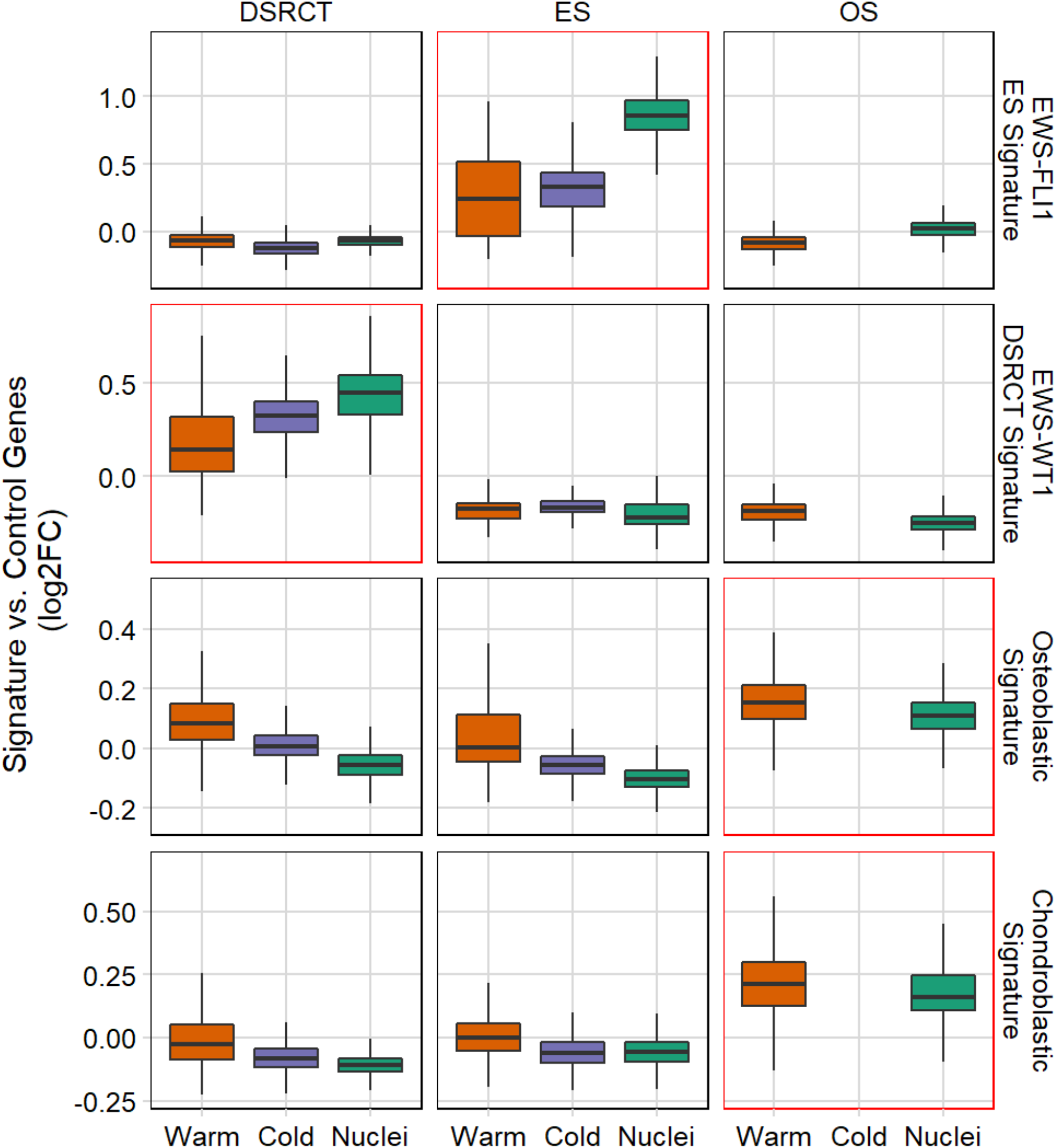
Biases associated with protocol on Sarcoma specific signatures. Sarcoma specific signatures were only detected in the respective sarcoma subtypes (red box). EWS-FLI1 ES signature is derived from a gene set of EWS-FLI1 target genes. EWS-WT1 DSRCT signature is derived from a gene set of EWS-WT1 target genes. Both osteoblastic and chondroblastic signatures are gene sets derived from reference databases.

### Single-nucleus RNA sequencing enriches for genes with long transcripts

Subsequent analysis revealed that several enriched genes in the Nuclei protocol are coded by transcripts longer in length compared to those enriched in the Warm protocol. To further investigate this interesting finding, we compared the gene lengths of commonly enriched genes in Warm versus Nuclei protocol for all sarcomas. We found that genes enriched in the Nuclei protocol had significantly longer genes (Wilcoxon test, p-value < 2.2e-16) (**Fig. S4A**).

This suggests that there is a possible gene length-associated bias in snRNA-seq. Recent work indicated that hybridization of the polyT RT-primer to intronic polyA stretches of nascent transcripts results in the gene length bias^17^. In fact, our analysis showed that 52% of reads for Nuclei mapped to intronic regions whereas 23% of reads were mapped for Warm protocol (**Fig. 6A**). We binned the genes into quartiles based on the gene length termed as Short (0 – 8077 nt), Short Med. (8077 – 24399 nt), Long. Med. (24399 – 66502 nt), and Long (> 66502 nt). On average, 55.6% of total genes greater than 66,502 nt (Long) were enriched in the Nuclei protocol compared to 28.2% and 26.6% in the Warm and Cold protocols, respectively (**Fig. 6B**). Interestingly, we also observed an opposite effect in the short genes (0 – 8077 nt) with 20.7% and 21.4% of the total genes in Warm and Cold protocols respectively as opposed to 4.1 % in the Nuclei protocol (**Fig. 6B**).

**Figure 6.**
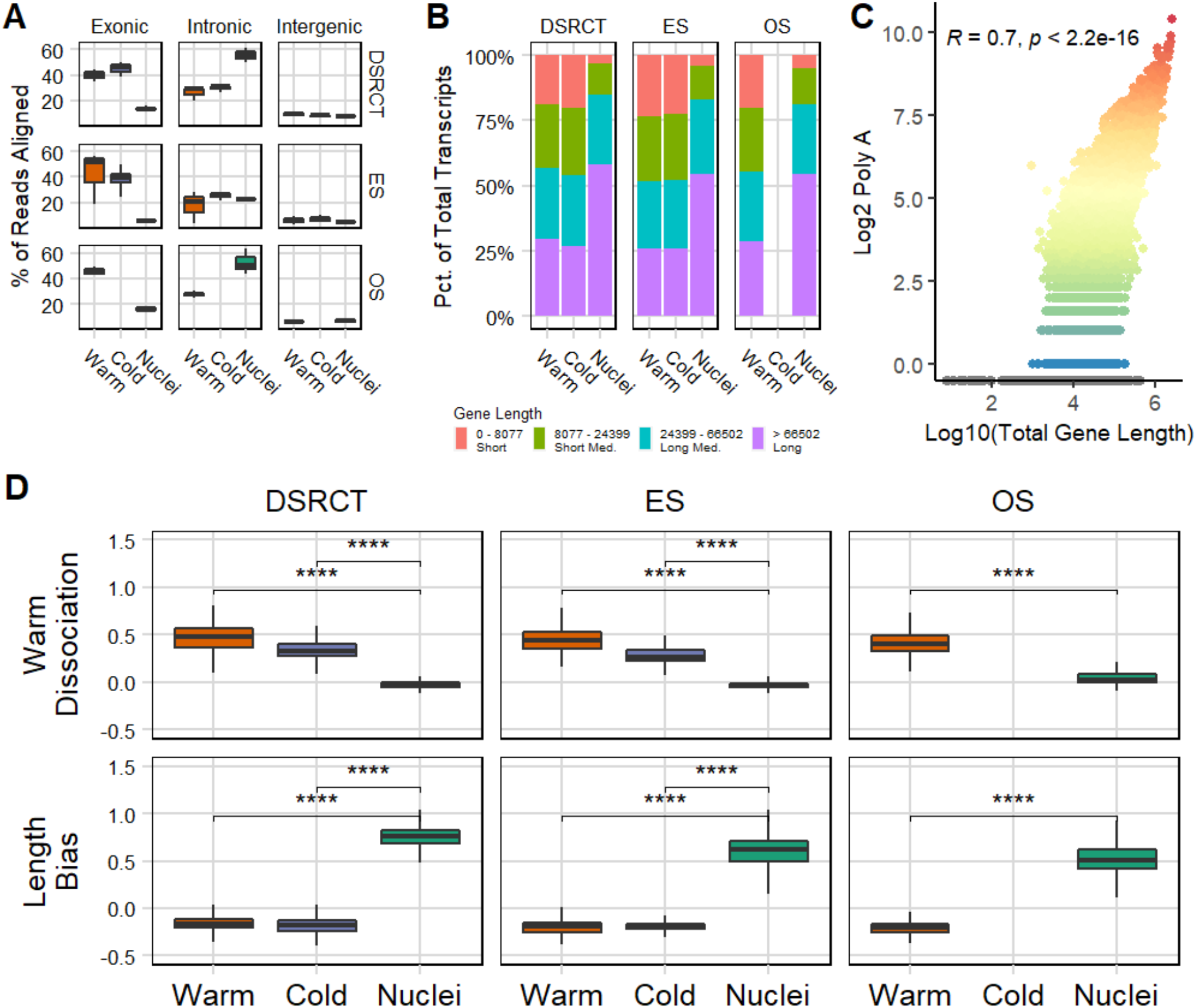
Gene-length associated bias in snRNA-seq. **a** Comparison of where transcriptomic reads are aligned. **b** snRNA-seq samples were enriched in genes with longer gene length. Genes were split into quartiles based on gene length. **c** Increased PolyA regions were associated with longer gene length. **d** Warm Dissociation score was significantly higher in the Warm and Cold protocol as opposed to the Nuclei protocol. Length Bias score was significantly higher in the Nuclei protocol (Wilcoxon test, **** denotes p <= 0.0001)

To answer if there were indeed a bias due to polyA stretches, we counted the number of polyA regions, defined as greater than 15 A repeats, within the full-length cDNA including both intronic and exonic regions for every gene. We observed a significant correlation between increasing gene length and polyA regions (R = 0.7, p-value < 2.2e16, (**Fig. 6C**). While we saw a positive correlation between scRNA and snRNA expression for each sarcoma type, there is a skew toward higher expression of genes that are longer and containing many polyA regions for snRNA data (**Fig. S4B-D)**.

### A gene length bias score accurately identified cells profiled by snRNA-seq

To evaluate the enrichment of long transcripts, we generated a length bias score by taking the top 200 genes with the highest number of polyA regions and combining them into a signature gene set (**Table S7**). Note that these genes were chosen purely by length and is agnostic to the underlying biology. In addition, we evaluated the expression of a previously generated warm dissociation signature from O’Flanagan et al.^4^ Our results demonstrated that the warm dissociation signature is clearly associated with the Warm protocol (**Fig. 5D**). On the other hand, expression of the length bias score is only observed in samples profiled using snRNA-seq. Together, these signatures robustly delineated the biases imparted by scRNA- and snRNA-seq for the different sarcoma subtypes.

To further illustrate this, we evaluated logistic regression models using the length bias and warm dissociation signatures to classify affected cells. We randomly split the ES data set into training and test groups. Using the logistic regression model, we could accurately predict samples that underwent the Nuclei protocol (AUC = 1.00) and whole-cells that displayed stress from the Warm protocol (AUC = 0.92) (**Fig. S5**). We applied the same model to the OS and DSRCT data set and observed the same findings (**Fig. S6**). To test if we could extrapolate this classifier to single-cell and single-nucleus libraries processed outside our lab, we used data from a recent paper^6^. In this work, the authors used collagenase type 4 at 37°C to dissociate a neuroblastoma PDX (O-PDX) into single cells and Tween with salts and Tris to dissociate O-PDX into single nuclei^6^. The authors demonstrated that the same dissociation signature we evaluated was elevated in whole-cells when compared to nuclei. Like our data, we applied the length bias and warm dissociation signatures to the O-PDX data set and were able to accurately predict samples that were nuclei (AUC = 1). Importantly, we observed the same findings in a neuroblastoma resection from a patient specimen to rule out if this was limited to only PDX specimens. Our results suggest that single nuclei could readily be classified just by the length bias score and that this may not be limited to just PDX samples (**Fig. S7)**. This implies that snRNA-seq enriches for longer transcript when compared to scRNA-seq from paired samples.

We suspected that when comparing sarcoma signatures between protocols for ES, the elevated expression of the EWS-FLI1 gene set in the Nuclei protocol may be due to the gene length-associated bias in snRNA-seq (**Fig. 4**). For each sarcoma signature, we divided the genes into four bins based on quartiles of gene length (**Fig. S8A**). As we expected, for the EWS-FLI1 target gene set, over 40% of genes in this specific set were considered long (> 66502 nt). This enrichment was not observed in the other gene sets. To further explore the effect of longer genes, we split the EWS-FLI1 target gene set into two groups – short (< 65502 nt) and long (> 65502 nt) genes. Next, we evaluated the resulting expression in ES and indeed observed a gene length-associated bias but only in the group that included long genes (**Fig. S8B**).

### Data integration recovers conserved markers and matching cell-states

As demonstrated by our UMAP embedding for OS (**Fig. 7**), the same samples processed simultaneously by scRNA-seq and snRNA-seq exhibit large batch effects and vastly different transcriptomic signatures. This complicates downstream analyses – even within the same cancer type – and will present unique challenges when investigators try to apply lessons learned from a dataset assessed by scRNA-seq to another generated in parallel using snRNA-seq.

**Figure 7.**
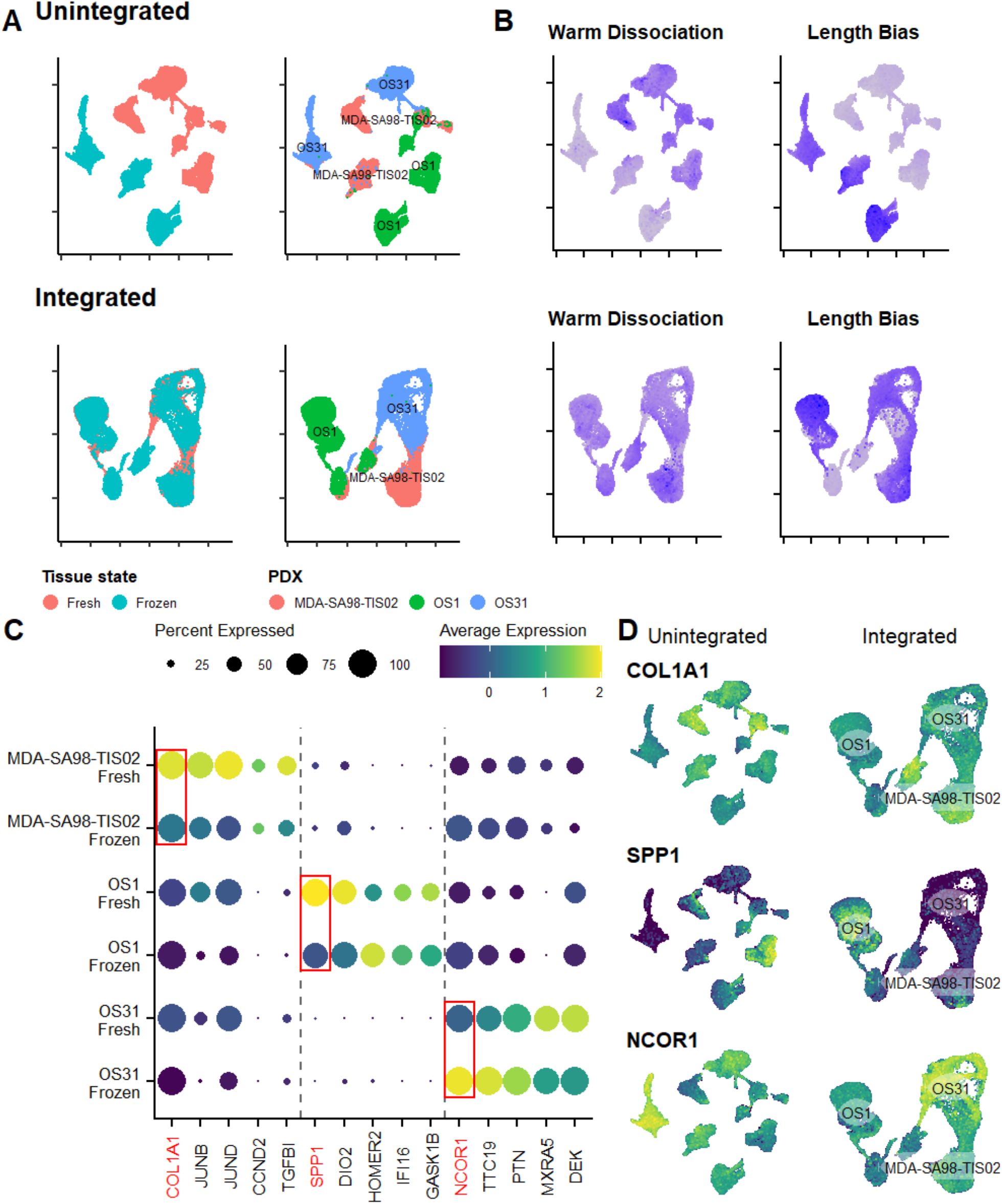
Integration recovers matching cell states from different dissociation methods for OS. **a** UMAP embeddings after integration (bottom row) showed alignment of matching PDX specimens. **b** After integration (bottom row), clusters on the UMAP were no longer affected by the identified biases. Darker blue indicates higher expression. **c** Dot plot of conserved markers show unique gene expression for each PDX. Top marker is in red. **d** After integration (right column), top conserved markers correctly aligned with the associated PDX cluster.

One could theoretically solve the dilemma using a brute-force approach that runs each sample twice, first by scRNA-seq then again using snRNA-seq, assuming of course that sufficient tissue exists, However, this method is likely to be impractical given tissue scarcity, frequent lack of paired fresh/frozen tissues, and the redundant costs associated with creating and sequencing the cDNA library twice per sample. Prospective collection of fresh tissues for rare cancers like sarcoma, or even rarer sarcoma subtypes like OS, ES, or DSRCT, presents additional hurdles.

To counter batch effects induced by sample handling, inter-operator variation, and differing technologies (e.g., CEL-Seq, Fluidigm, 10x Chromium), several bioinformatic tools exist to remove covariates. Utilizing the integration workflow for Seurat v3, we corrected for protocol biases and integrated each dataset of matched PDX specimens^18^. Each sarcoma subtype dataset was split by dissociation method and then integrated.

As can be seen after applying integration, technical biases from dissociation protocols are mitigated, and PDX specimens with similar cell states now cluster together (**Fig. 7A**). As an example, the Warm dissociation signature that was previously enriched in fresh tissues analyzed by scRNA-seq, is now homogeneously distributed and not affecting the clustering of samples (**Fig. 7B**). Similarly, the length bias score, which was previously causing the samples profiled by snRNA-seq to separate is now mitigated (**Fig. 7B**).

When observing conserved markers for OS PDXs, we predicted that both fresh and frozen from the same PDX would exhibit the same DEGs when compared to other PDXs. Based on the Figure 6C, we observed that the DEGs are conserved between fresh and frozen specimens. As we expected, the unintegrated UMAP does not neatly align the conserved markers, which is most likely due to the technical biases that we have shown influencing the algorithm (**Fig. 7D**, left column). Upon integration of the samples, we can clearly observe the conserved DEGs localizing to each PDX (**Fig. 7D**, right column). We extended this same analysis to both ES and DSRCT, and we observed the same effects (**Fig. S9, 10**). Like batch corrections, this illustrates the need for properly integrating diverse datasets, where in this work paired specimens underwent different dissociation protocols, to reliably perform downstream analyses.

## Discussion

The advent of single-cell transcriptomic profiling has revolutionized the ability to decipher gene expression in a way that would have been otherwise unimaginable just a decade ago. Of major value for cancer research is the opportunity to measure cellular composition of each tumor, as well as the individual states and phenotypes of individual cancer cells that would have otherwise been obscured with whole-tumor RNA-seq approaches. Accurate interpretation of the results, however, requires a keen appreciation for the technical and computational biases introduced by the chosen methods for tissue handling, cell dissociation, and cell or nuclei preservation.

In this work, we sought to elucidate the inherent biases of different dissociation protocols on the transcriptome of sarcomas, focusing initially on three subtypes that predominantly affect children and young adults. To avoid consuming scarce clinical research specimens, we limited our research scope to early-generation sarcoma PDXs, which maintain close fidelity to the OS, ES, or DSRCT patients from whom they were derived. The choice to use PDXs, rather than human tumors, also stemmed from our ability to tightly regulate how tumors were collected, stored, and processed. Further, the PDX tissues afforded an opportunity to receive fresh and snap frozen tissue simultaneously from the same tumor to avoid temporal biases. In contrast, the human tumors as they exist in our institution, were collected months or years apart, often at different points in each patient’s treatment course, and typically snap frozen or formalin-fixed and paraffin-embedded without gathering a fresh tissue comparator.

Our work builds upon prior studies in normal tissues and carcinomas that have analyzed the protocol dependent biases used for scRNA-seq and snRNA-seq^4-6,18-20^. Consistent with prior studies, enzymatic digestion at 37°C invoked a marked stress-response, manifest by upregulation of immediate early genes (IEGs), such as FOS, JUN, MYC^4^. As expected, this stress-response was minimized in the Cold protocol and almost absent in the Nuclei dissociation.

Interestingly, because many sarcoma subtypes are caused by chromosomal translocations that produce pathognomonic fusion proteins, we had the opportunity to determine if protocol-specific technical biases interfered with the downstream target gene signatures induced by EWS-FLI1 or EWS-WT1 in ES and DSRCT, respectively. Though we hypothesized a stress-response could affect the expression of EWS-FLI1 target genes, we observed in fact that snRNA-seq had a significantly greater impact, possibly due to enrichment for genes with longer transcripts. This unexpected bias towards longer transcripts resulted in an EWS-FLI1 target gene set that was overexpressed in samples assessed by snRNA-seq, as opposed to scRNA-seq.

As to why the EWS-FLI1 target genes contain an overabundance of long genes, we explored a few possibilities. The EWS-FLI1 transcription factor is known to bind to GGAA microsatellite repeats of 9 or more^21,22^. This may be influenced with transcription length like the increase of polyA region with increasing length. However, many of the microsatellite repeats that enable EWS-FLI1 binding were found within the first intron or the promoter region, which could be located as far as 1 Mb upstream of the transcription start sites^22^. A more likely explanation may be found in the broader analysis of long genes. A review of the effect of gene length found positive correlations with intron number, protein size, and SNPs^23^. Remarkably, gene length is also associated with cancer, heart diseases, and neuronal development^23,24^. Given that a portion of EWS-FLI1 targets are known to be neural genes, we can speculate that some of the long genes in the EWS-FLI1 gene set are neural related^25^.

Overall, special care must be considered when comparing data between whole-cells and nucleus. To remove the technical bias introduced by snRNA-seq, we generated a length bias signature using genes with long transcripts. Others have shown that technical biases or batch-to-batch effects can be regressed from snRNA- or scRNA-seq data^19^. Regression of the length bias from the snRNA-seq can produce comparable results to scRNA-seq^17^. However, comparing whole-cell and nucleus transcriptomes between specimens of different tissue origin or disease should be interpreted with caution. As noted already, gene length is associated with cancer, heart diseases, and neuronal development and correlated with SNPs^23,24^. On the other hand, our data and others have demonstrated that snRNA-seq data is enriched with lncRNA as compared to scRNA-seq^13^. While this may seem like a confounding variable when trying to compare the two different modalities (i.e., scRNA- and snRNA-seq), it may be beneficial to utilize snRNA-seq if the intent is to enrich and study lncRNA that regulate cell biology.

Computational methods play an important role normalizing data for known technical biases. After applying Seurat v3 integration, matched PDX specimens with similar cell states clustered together on the UMAP embedding. This is to be expected since Seurat v3 jointly reduces the dimensionality of datasets using a diagonalized CCA to identify shared biological markers and conserved gene expression signatures^18^. The algorithm then finds mutual nearest neighbors in this low-dimensional representation to recover matching cell states between datasets^26^. Since feature selection for integration is limited to variable features within each dissociation protocol, subtle differences between protocols (such as the warm dissociation signature) will play a smaller role.

Not performed in this study but an important concept to highlight when using different dissociation protocols is the effect on cellular composition bias. While scRNA-seq and snRNA-seq adequately represent the original cell populations, others have noted some differences, especially for immune cells^6,7^. An unavoidable limitation of our study was the placement of PDXs within immunocompromised murine models that lack a full immune cell repertoire. Thus, we did not have the opportunity to assess whether snRNA-seq underestimates the prevalence of T-cells, B-cells, and NK cells, as has been reported previously in carcinomas^27^. Others have shown the methanol fixation was superior to cryopreservation with respect to epithelial cell preservation, and it remains to be explored whether one preservation method is superior to another in retaining the native cell distribution or sarcomas or normal mesenchymal tissue. As spatial image omics (SIO) gains traction, one could envision using this technology as a ‘gold-standard’ to meticulously catalog a cancer’s true cell composition without suffering the aforementioned technological artifacts^28^.

Our work is the first to rigorously compare the protocols used for sc/snRNA-seq to assess their effect on gene expression in sarcoma tissues. Consistent with prior reports in epithelial malignancies, we demonstrate that Warm dissociation introduced a similar cell stress signatures in three pediatric sarcoma subtypes. Among other key findings, the gene signatures associated with ES’s and DSRCT’s fusion proteins were more readily observed using snRNA-seq. This result has immediate relevance, since it suggests that pre-existing frozen specimens can be used to advance sarcoma research. Last, we demonstrate that computational algorithms can be used to remove some of the biases linked to the experimental methods.

## Materials and Methods

### Collection of fresh tissue for scRNA-seq

All experiments were conducted per protocols and conditions approved by the University of Texas MD Anderson Cancer Center (MDACC; Houston, TX) Institutional Animal Care and Use Committee (eACUF Protocols #00000712-RN02). Male NOD (SCID)-IL-2Rg^null^ mice (The Jackson Laboratory; Farmington, CT) were subcutaneously injected with PDX explants (2 mm) to generate xenografts. All mice were maintained under barrier conditions and treated using protocols approved by The University of Texas MD Anderson Cancer Center’s Institutional Animal Care and Use Committee. MDA-SA98-TIS02, OS1, and OS31, are PDX lines maintained by the Pediatric Solid Tumors Comprehensive Data Resource Core^29^. Once their tumors reached a volume of 150 mm^3^, tumors were explanted and a portion was flash-frozen for snRNA-seq, while the remainder underwent dissociation.

### Dissociation workflow from fresh solid tumor samples

Samples were collected and immediately placed into MACS® Tissue Storage Solution (Miltenyi Biotec) and kept on ice during transport. On arrival to the laboratory, samples were minced using a scalpel into fragments under ∼ 0.5 mm under aseptic conditions. Next, samples were evenly split for either warm or cold enzymatic dissociation.

For warm dissociation of ES and DSRCT PDX specimens, the human Tumor Dissociation Kit (Miltenyi Biotec) was used. The dissociation was performed under manufacturer’s protocol using the gentleMACS™ Dissociator (Miltenyi Biotec), a benchtop instrument for the semi-automated dissociation of tissues into single-cell suspensions. The gentleMACS Program sequenced followed the suggestion for ‘Soft’ Tumor type. Following completion of the program, 2x volume of media was added to the samples. This was followed by filtration through a MACS SmartStrainer (70 μm, Miltenyi Biotec) and centrifugation at 300g for 5 min. Cells were resuspended in 90% FBS and 10% DMSO at a concentration of 1 million cells per mL and placed in a Thermo Scientific™ Mr. Frosty™ Freezing Container in a -80 °C freezer.

For warm dissociation of OS PDX specimens, tissue was minced into 3 – 4 mm pieces with sterile a scalpel or scissor. The tissues were washed several times with Hank’s Balanced Salt Solution (HBSS). HBSS was next aspirated and dissociation buffer (HBSS, 1 mg/mL collagenase, 3mM CaCl_2_, 1 μg/mL DNase) was added to submerge the tissue. Tissue is then incubated at 37 °C for up to 12 hours. The cell suspension was then filtered using a 40 μm cell strainer. The filtrate is pelleted using a cenfriifugation at 400g for 5 min. Cells were resuspended freezing meddium and placed in a Thermo Scientific™ Mr. Frosty™ Freezing Container in a -80 °C freezer.

For cold dissociation, protocol was adapted from *Adam et al*^5^. Cold protease solution was prepared from 5 mM CaCl_2_, 10 mg/mL *B. Licheniformis* protease, and 125 U/mL DNase I in 1x PBS. Tissue was minced using a scalpel into fragments under 0.5 mm. Fragments were placed in a MACS C-tube and 5 mL of ice-cold cold protease solution was added. The samples were incubated for 10 min at 4 °C with rocking. This was followed by placing the samples in a gentleMACS™ Dissociator (Miltenyi Biotec) and running the m_brain_03 program twice in succession. Afterwards, the samples were centrifuged at 300g for 5 min and resuspended in 3 mL of trypsin-EDTA for 1 min at room temperature. The trypsin-EDTA was then neutralized using ice-cold 10% FBS in 1x PBS and triturated. This was followed by filtration through a MACS SmartStrainer (70 μm, Miltenyi Biotec) and centrifugation at 300g for 5 min. Cells were resuspended freezing medium at a concentration of 1 million cells per mL and placed in a Thermo Scientific™ Mr. Frosty™ Freezing Container in a -80 °C freezer. Cryovials were moved to LN2 storage for long-term.

### Thawing cryopreserved cells

The cells were removed from LN_2_ or -80 °C freezer, if they were reccently cryopreserved, and placed into a 37 °C water bath for 3 min. The contents were then transferred to a 15 mL centrifuge tube. 1 mL of complete medium was used to wash the cryovial and added drop-wise into the centrifuge tube. Next, 8 mL of complete medium was added drop-wise to reduce osmotic shock. Cells were then centrifuged at 300g for 5 min and resupsneded in 1x PBS supplemented with 0.04% BSA. This was followed by live cell enrichment using FACS. Single-cell suspensions were stained with Caclein AM live cell stain and SYTOX™ Red dead cell stain.

### Nuclei Isolation workflow

The protocol was adapted from Habib *et al*^9^. We isolated nuclei from fresh-frozen tissue using the Nuclei EZ Prep Kit (Sigma-Aldrich). Fresh-frozen tissue specimen were cut into piecces < 5 mm over dry ice and then placed in 0.5 mL ice-cold EZ lysis buffer. This was followed by homogenizing using a Chemglass Life SciencesSupplier BioVortexer Mixer (Fisher Scientific) attached with a plastic microcentrifuge pestle on ice. Then 1 mL of ice-cold EZ lysis buffer was added and samples were incubated on ice for 5 min. Debris was filtered out using a pluriStrainer Mini 70 μm into a new tube. This was followed by centrifugation at 500g for 5 min. Samples were then incubated with 1 mL of ice-cold EZ lysis buffer on ice for 5 min followed by centrifugation. Afterwards, the supernatant was aspirated and 0.5 mL of Nuclei Wash and Resuspension Buffer (NWRB, 1X PBS supplemented with 1.0 % BSA and 0.2U/μl RNase Inhibitor) was carefully added without disrupting the pellet, which was followed by 5 min of incubation. Next, we added 0.5 mL of NWRB and centrifuged at 500g for 5 min. We repeated the wash and incubation once more, followed by centrifugation. The supernatant was aspirated and the nuclei were resuspended in NWRB. A portion was visualized with Trypan blue under the microscope to inspect for debris and nuclei integrity.

To sort nuclei, single-nucleus suspensions were stained with 7-AAD in NWRB for 5 min on ice. Then a BD cell sorter was used to sort up to 100,000 7-AAD positive events. Quality control of post-sort nuclei concentration was evaluated under a microscope to ensure adequate count. This was followed by loading nuclei onto a 10X chip.

### Library preparation and sequencing

We followed the standard protocol set by 10x Genomics for single-cell/single-nucleus capture. A targeted capture of 5000 single cell or single nucleus were loaded onto each channel of a Chromium single-cell 3’ Chip. The single cells and single nuclei were partitioned using the gel beads within the Chromium Controller. Afterwards, we performed cDNA amplification and fragmentation. This was followed by index and adapter attachment. Samples were pooled and sequenced on a NovaSeq 6000 with targeted sequence depth at 100,000 reads/cell or nucleus.

### sc/snRNA-seq data preprocessing

We used Cell Ranger mkfastq to generate demultiplexed FASTQ files. Reads were aligned to human GRCh38 genome and reads were then quantified as UMIs buy Cell Ranger count. For snRNA-seq, reads were mapped with both introns and exons in Cell Ranger 5.0 using the include-introns option for counting intronic reads^10^.

We performed QC and normalization separately for each sarcoma PDX. We followed the guidelines for QC from OSCA and others^20^. We inspected UMIs, gene counts, and percentage of mitochondrial genes and identified outliers based on median absolute deviation (MAD). We used a strict value of 2 or more MADs from the median while also using generic cut offs. Cells that did not meet the criteria were removed from analysis. Scrublet was used to predict and detect doublets within the data^30^. While doublets were flagged, there was not a single cluster of doublets, which would be evident as an artifact, so no cells were removed.

### Data normalization, dimensional reduction, and comparisons

Seurat v3 was used for sample normalization, dimensional reduction, scaling, and differential expression analysis^18^. We used Wilcoxon test to compare gene expression between protocols. Enrichr was used for pathway enrichment. We set a log2 fold change threshold of log2(1.5) or greater. This will result in genes that are 50% greater than the baseline. The AddModuleScore function in Seurat v3 was used to observe the averaged gene expression of different gene set. We used curated gene sets of a warm dissociation signature from O’Flanagan et al.^4^, EWS-FLI1 gene targets^15^, EWS-WT1 gene targets^16^ and osteoblastic and chondroblastic signatures classically associated with the tissue origin of OS (**Table S6**). The osteoblastic and chondroblastic signatures were found on Harmonizome (https://maayanlab.cloud/Harmonizome/). The osteoblastic signature was specifically found in the GeneRIF Biological Term Annotations under ‘Osteoblastic’. The chondroblastic signature was specifically found in the TISSUES Text-mining Tissue Protein Expression Evidence Scores under ‘Chondroblasts’. To find conserved markers between dissociation methods, we used the function FindConservedMarkers in Seurat v3. We performed integration using the integration functions within Seurat v3. The datasets were integrated by dissociation protocol.

### Predicting sample type by bias scores

To classify nuclei and cell using the length bias and warm dissociation scores, data sets were randomly split into a training and test set. To prevent data leakage, scaled data was not used. We then calculated the gene set scores separately on the training and test sets. A logistic regression model was fit to the training set on either the warm dissociation or length bias score to predict for cells and nuclei respectively. We calculated the probabilities and the area under the curve using the pROC v1.18.0 package. This was compared to a random gene signature equal in number of genes of either length bias or warm dissociation gene sets.

## Declarations

### Ethics Approval

All experiments were conducted per protocols and conditions approved by the University of Texas MD Anderson Cancer Center (MDACC; Houston, TX) Institutional Animal Care and Use Committee (eACUF Protocols #00000712-RN02).

### Consent for Publication

N/A

### Availability of data and materials

The datasets generated analyzed in this study are available at the Gene Expression Omnibus (GEO, https://www.ncbi.nlm.nih.gov/geo/) repository accession no. GSE200529. Additional datasets analyzed in this study can be found at GSE140819.

### Authors’ contributions

DDT contributed to the conception, design, analysis, and interpretation of data. SC, RWP, SK, and JS contributed to sample acquisition and dissociation. RG, SC, NG, NCD, VG, and AG contributed to specimen database and clinical information. VG, NG, NCD contributed to interpretation and review of data. SC, AJL, and JA Livingston contributed conception and design. DDT and JA Ludwig contributed to the writing, review, and revision of the manuscript with input from all authors. JA Ludwig and SC contributed to study supervision. All authors read and approved the final manuscript.

## Acknowledgements

The authors acknowledge the support of the High-Performance Computing for research facility at the University of Texas MD Anderson Cancer Center for providing computational resources that have contributed to the research results reported in this paper. The authors acknowledge the support of the Cancer Prevention Research Institute of Texas (CPRIT) Pediatric Solid Tumors Comprehensive Data Resource Core (RP 180819) for providing tissue specimens and Energy Transfer Partners for the additional support provided to collect and store tissue specimens.

