## Supplementary Materials for "Dissociation Protocols used for Sarcoma Tissues Bias the Transcriptome observed in Single-cell and Single-nucleus RNA sequencing"

### Additional Figures

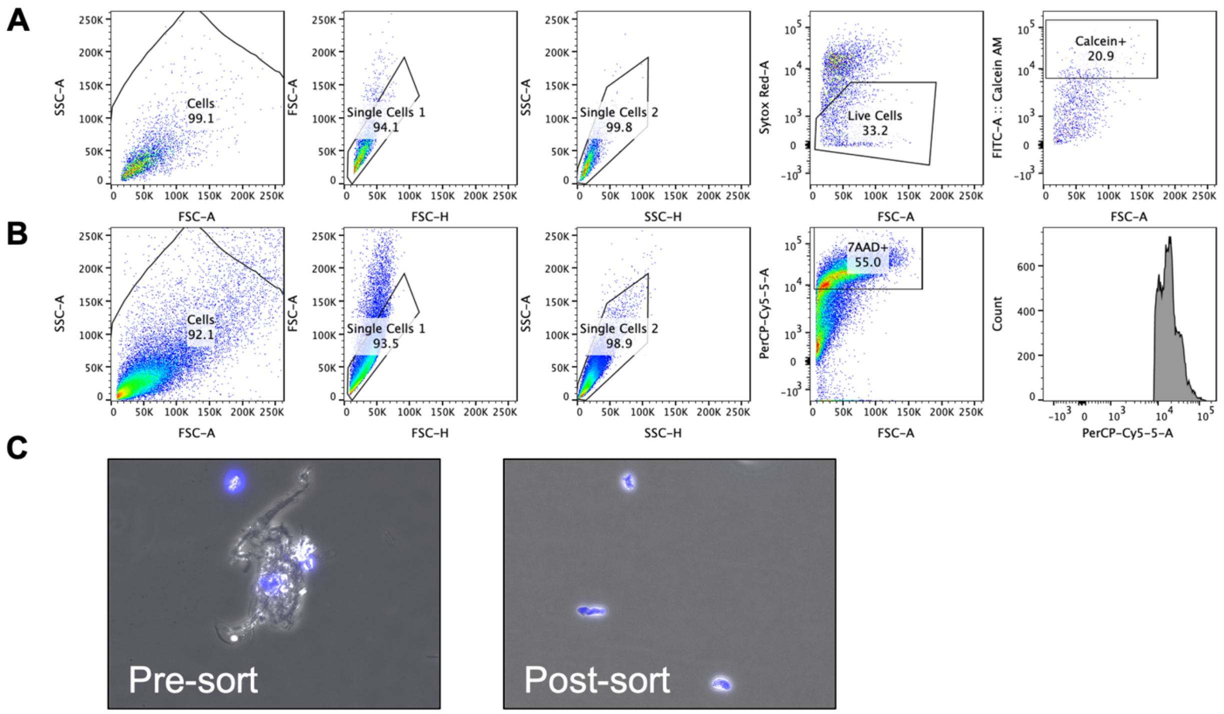


**Figure S1. Gating strategy to enrich for single-cells and single-nuclei. a** Cells are gated by SSC-A and FSC-A (first plot). Doublets are removed first by FSC-A and FSC-H and then by SSC-A and SSC-H (second and third plots). Live cells are gated from dead cells by Sytox Red-A (dead cells) and FSC-A (fourth plot). Live cells are sorted using FITC-A Calcein-AM (fifth plot). **b** Nuclei are gated by SSC-A and FSC-A (first plot). Doublets are removed first by FSC-A and FSC-H and then by SSC-A and SSC-H (second and third plots). Nuclei are gated from debris and intact cells by PerCP-Cy5-5-A (7-AAD, fourth plot). Histogram shows DNA content of nuclei (fifth plot). **c** Staining of nuclei with DAPI pre- and post-sort. Single nucleus without debris are enriched with sorting.

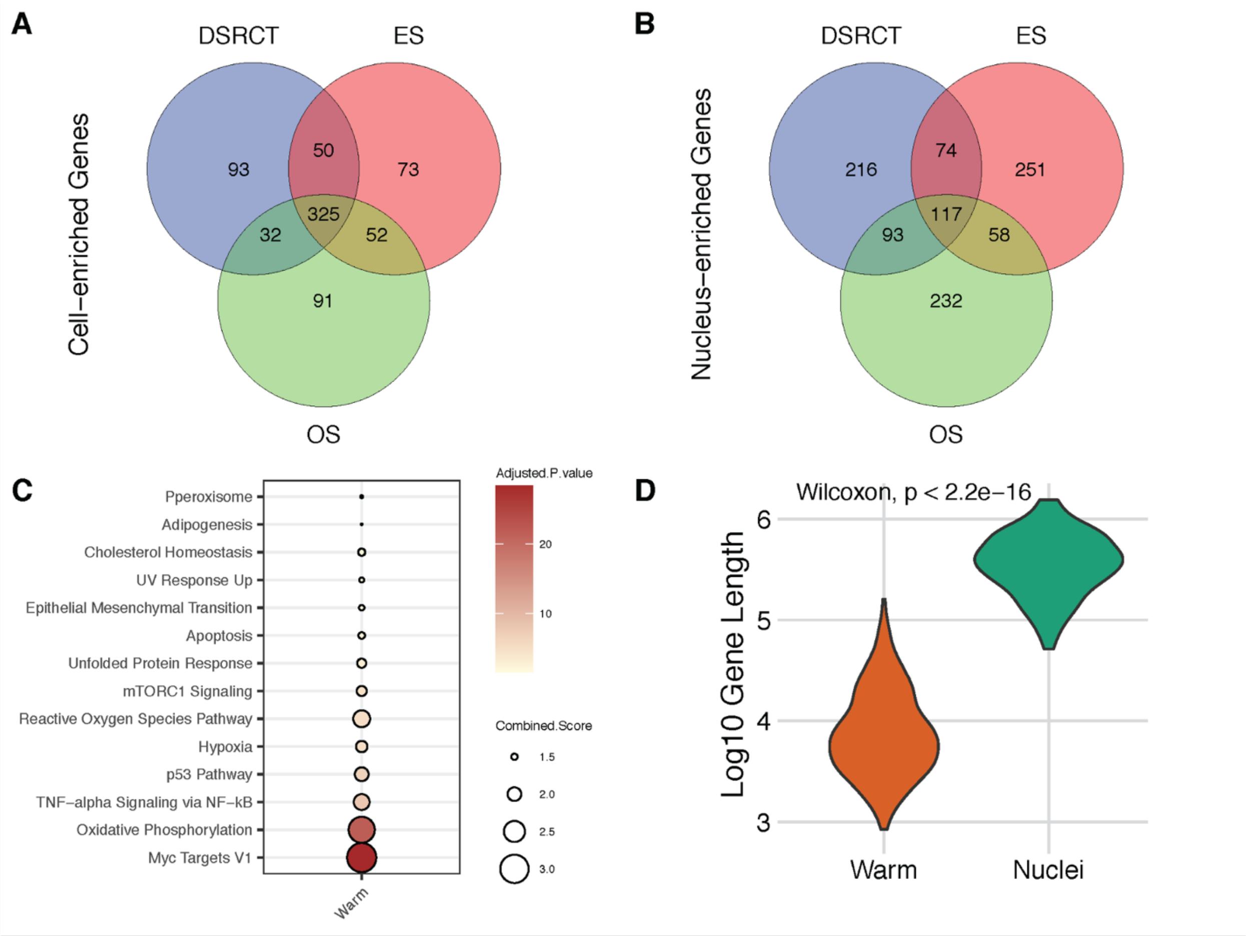


**Figure S2. Consensus Differentially Expressed Genes between Sarcoma Subtypes. a** Venn diagram of cell-enriched genes with 325 consensus genes enriched in the Warm protocol. **b** Venn diagram of nucleus-enriched genes with 117 consensus genes enriched in the Nuclei protocol. **c** Dot plot of enrichR scores of the Hallmark gene sets from MSigDB using only consensus genes. **d** Genes enriched in Nuclei were significantly longer than those in Warm ($p < 2.2e\text{-}16$).

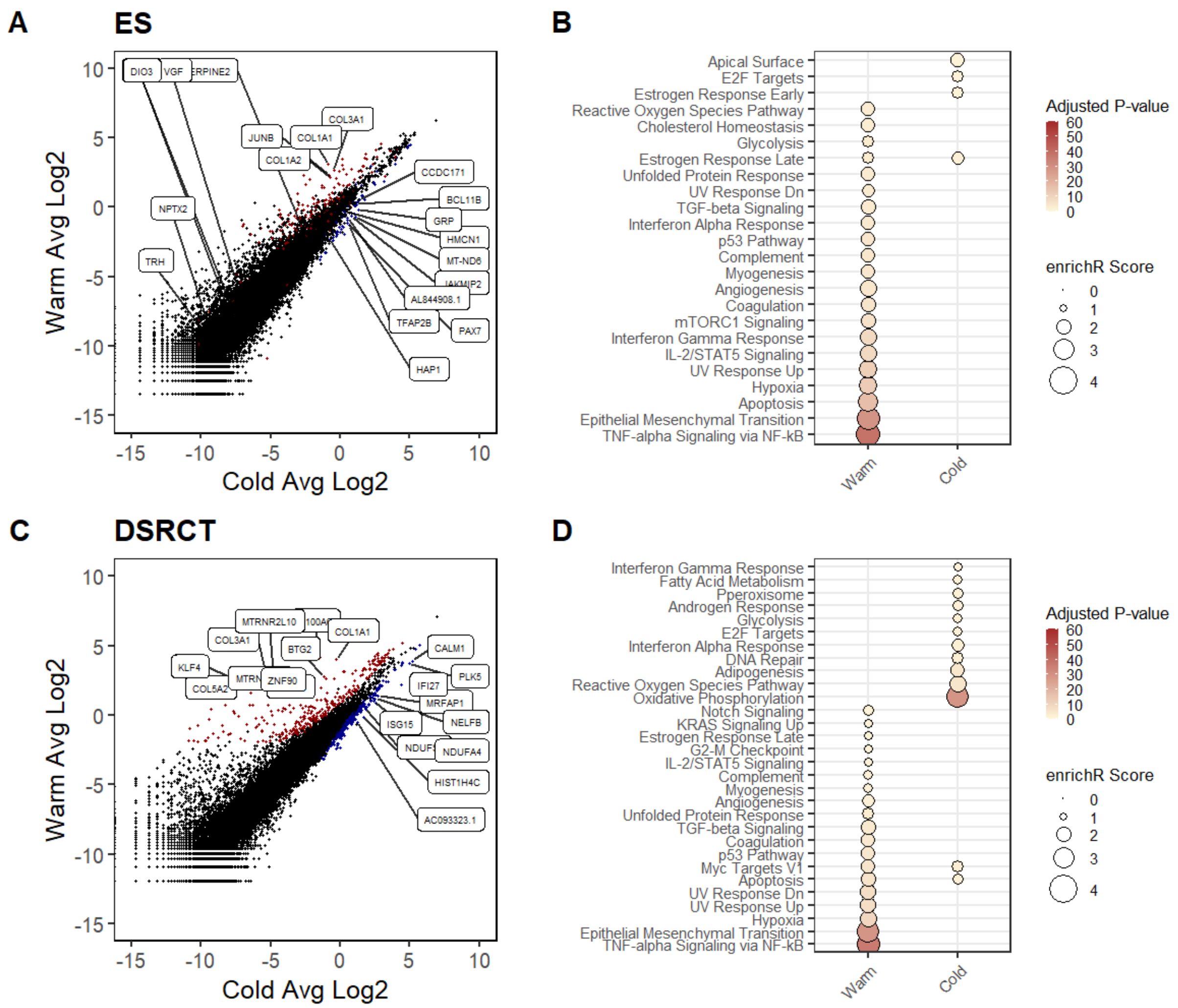


**Figure S3. DEGs Biases introduced by Warm and Nuclei Protocols.** Scatter plot of log transformed gene expression levels between Warm and Cold. Red indicates up-regulated in Warm, and blue indicates up-regulated in Cold with p-value < 0.05. Black is non-significant. Dot plot of enrichR scores of the Hallmark gene sets from MSigDB. Plots are shown for DSRCT **a, b**; and ES **c, d**.

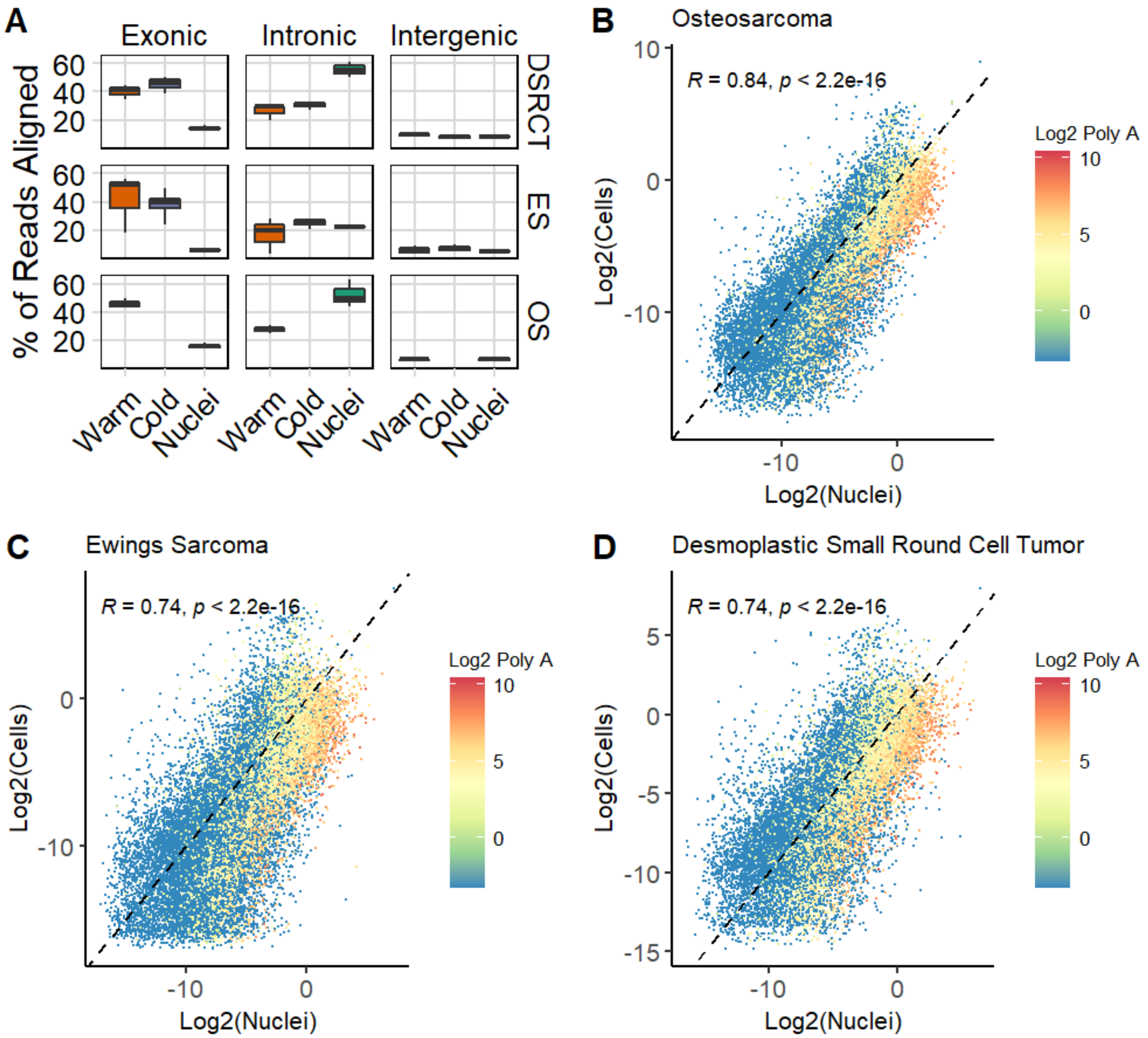


**Figure S4. Differences in Sequencing Performance. a** Comparison of where reads are aligned. Log-log plots of gene expression between cells and nuclei colored by number of PolyA regions for **b** OS, **c** ES, and **d** DSRCT.

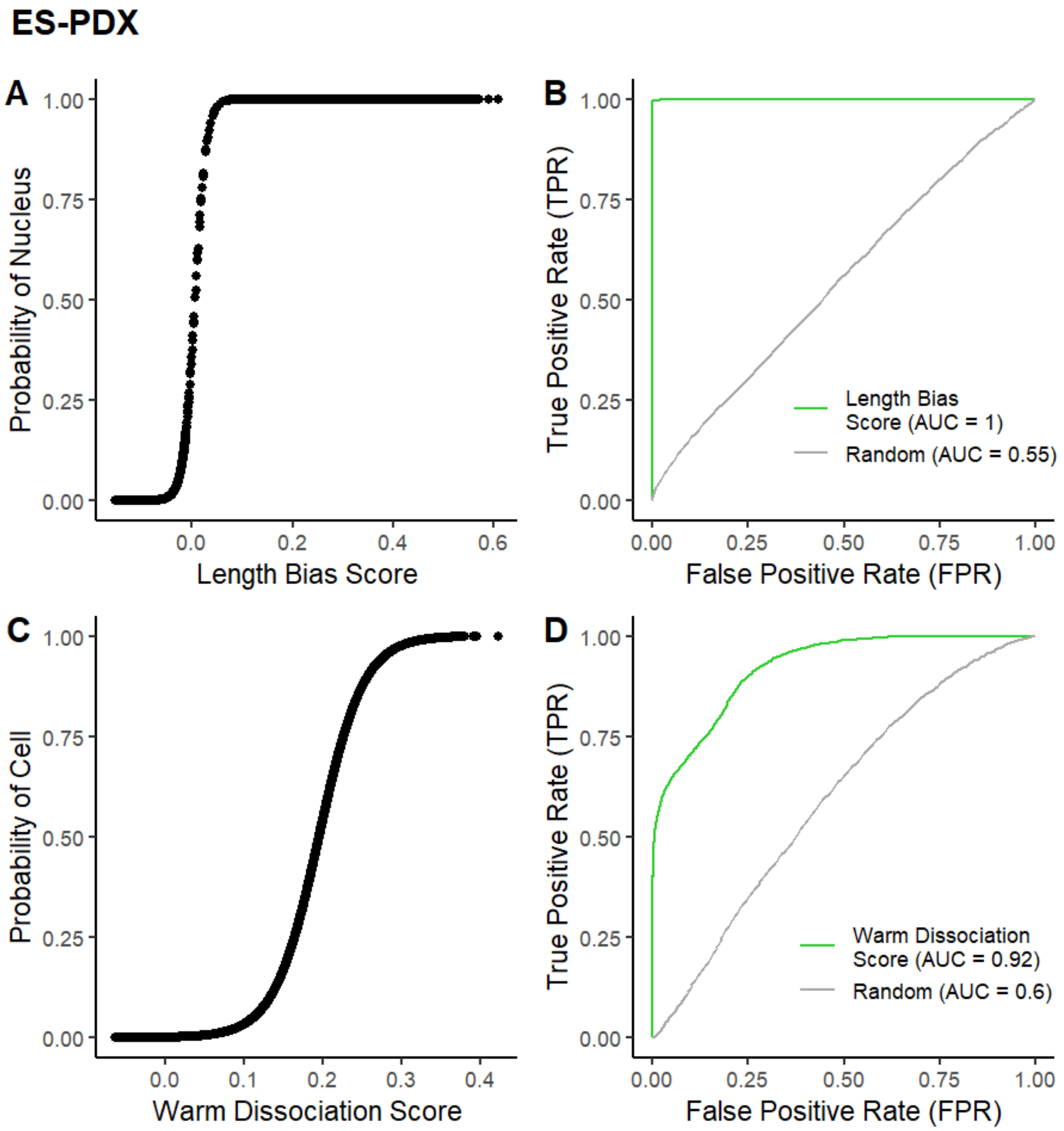


**Figure S5. Classifying sample type based on bias scores for ES. a** Predicting nuclei on test data after training data on Length Bias Score using a logistic regression. **b** Receiver operating characteristic (ROC) curve was used to evaluate the performance of the Length Bias Score to classify nuclei. **c** Predicting cells on test data after training data on Warm Dissociation Score using a logistic regression. **d** ROC curve was used to evaluate the performance of the Warm Dissociation Score to classify cells.

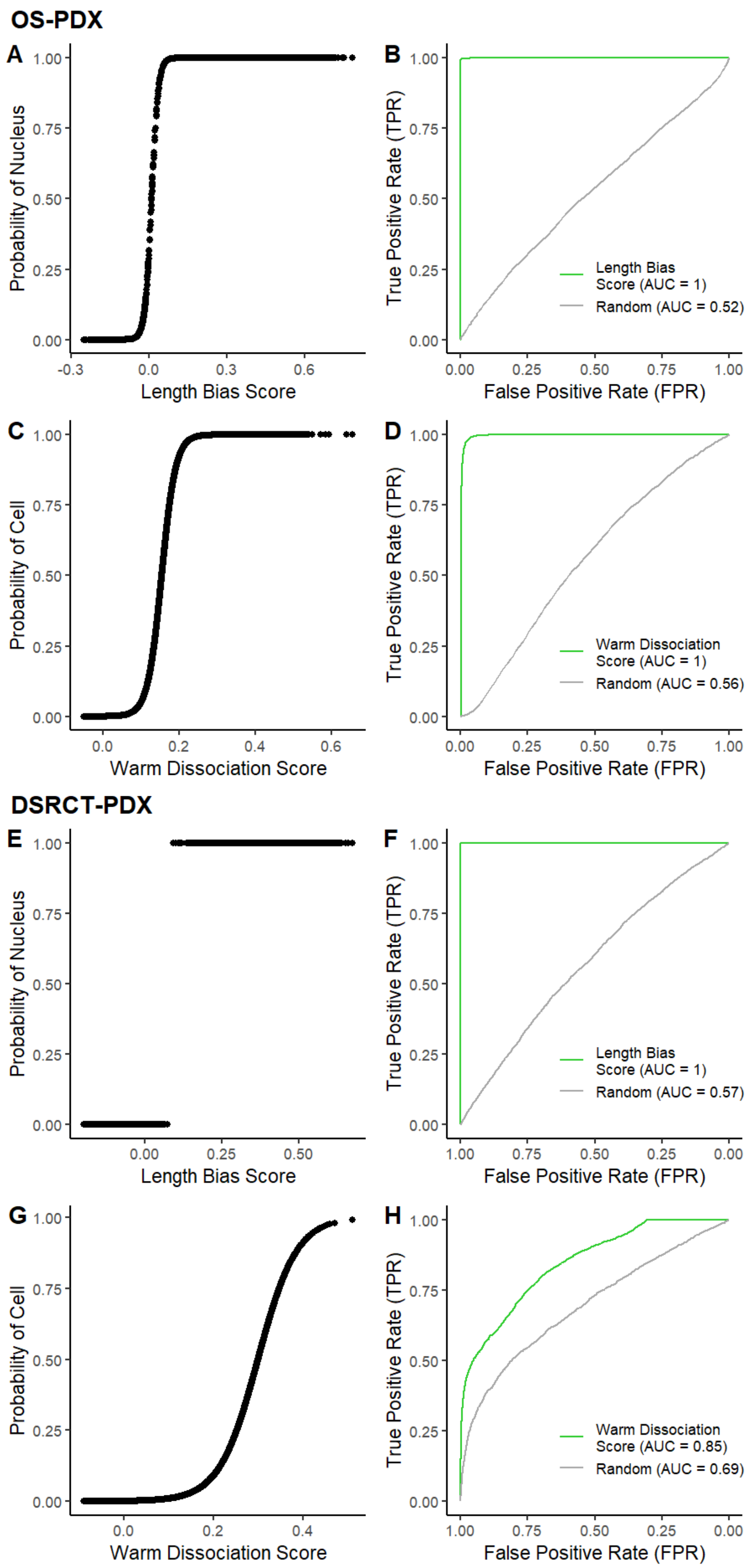
OS-PDX
A
Probability of Nucleus
1.00
0.75
0.50
0.25
0.00
-0.3
0.0
0.3
0.6
Length Bias Score
B
True Positive Rate (TPR)
False Positive Rate (FPR)
0.00
0.25
0.50
0.75
1.00
Length Bias
Score (AUC = 1)
Random (AUC = 0.52)
C
Probability of Cell
0.0
0.2
0.4
0.6
Warm Dissociation Score
D
Warm Dissociation
Score (AUC = 1)
Random (AUC = 0.56)
DSRCT-PDX
E
Probability of Nucleus
0.00
0.25
0.50
Length Bias Score
F
Length Bias
Score (AUC = 1)
Random (AUC = 0.57)
G
Probability of Cell
0.0
0.2
0.4
Warm Dissociation Score
H
Warm Dissociation
Score (AUC = 0.85)
Random (AUC = 0.69)

**Figure S6. Classifying sample type based on bias scores for OS and DSRCT. a** Probability of nuclei based on logistic regression model of Length Bias Score for OS. **b** ROC curve for using Length Bias Score to classify nuclei. **c** Probability of cell based on logistic regression model of Warm Dissociation Score for OS. **d** ROC curve for using Warm Dissociation Score to classify cells. **d** Probability of nuclei based on logistic regression model of Length Bias Score for DSRCT. **f** ROC curve for using Length Bias Score to classify nuclei. **g** Probability of cell based on logistic regression model of Warm Dissociation Score for DSRCT. **h** ROC curve for using Warm Dissociation Score to classify cells.

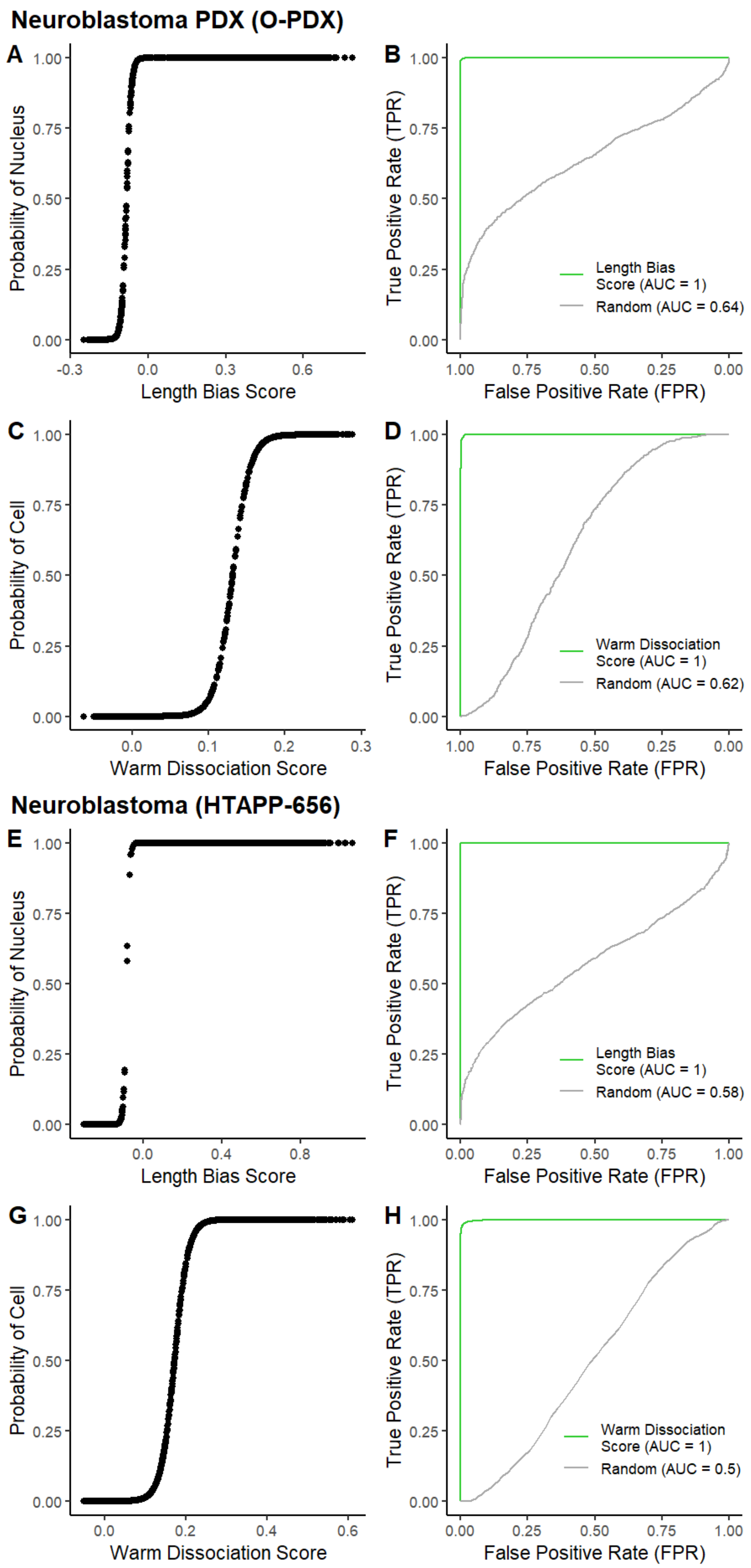

Neuroblastoma PDX (O-PDX)
A
Probability of Nucleus
Length Bias Score
B
True Positive Rate (TPR)
False Positive Rate (FPR)
Length Bias Score (AUC = 1)
Random (AUC = 0.64)
C
Probability of Cell
Warm Dissociation Score
D
True Positive Rate (TPR)
False Positive Rate (FPR)
Warm Dissociation Score (AUC = 1)
Random (AUC = 0.62)
Neuroblastoma (HTAPP-656)
E
Probability of Nucleus
Length Bias Score
F
True Positive Rate (TPR)
False Positive Rate (FPR)
Length Bias Score (AUC = 1)
Random (AUC = 0.58)
G
Probability of Cell
Warm Dissociation Score
H
True Positive Rate (TPR)
False Positive Rate (FPR)
Warm Dissociation Score (AUC = 1)
Random (AUC = 0.5)

**Figure S7. Classifying sample type based on bias scores for Neuroblastoma. a** Probability of nuclei based on logistic regression model of Length Bias Score for O-PDX. **b** ROC curve for using Length Bias Score to classify nuclei. **c** Probability of cell based on logistic regression model of Warm Dissociation Score for O-PDX O-PDX. **d** ROC curve for using Warm Dissociation Score to classify cells. **d** Probability of nuclei based on logistic regression model of Length Bias Score for HTAPP-656. **f** ROC curve for using Length Bias Score to classify nuclei. **g** Probability of cell based on logistic regression model of Warm Dissociation Score for HTAPP-656 HTAPP-656. **h** ROC curve for using Warm Dissociation Score to classify cells.

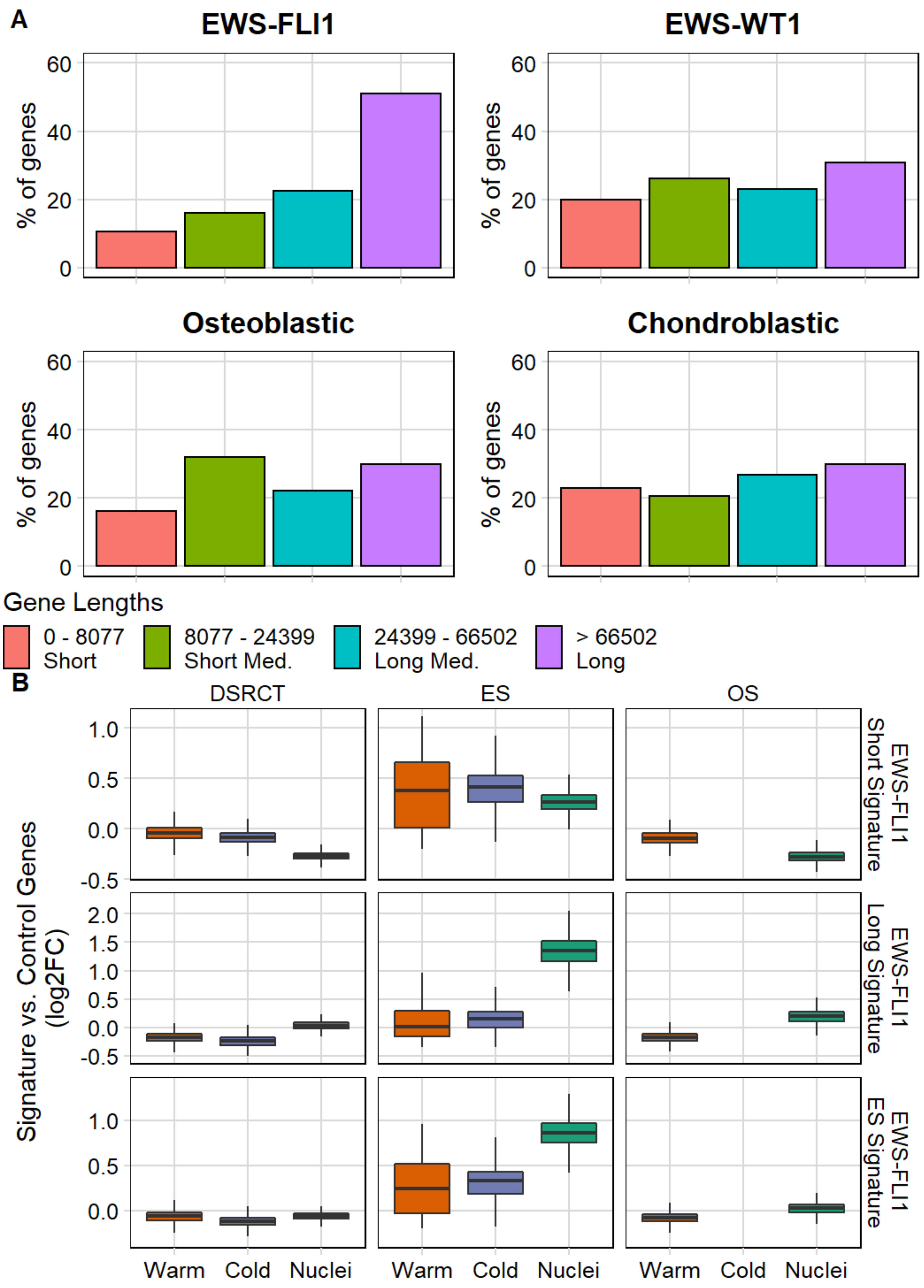
A
EWS-FLI1
EWS-WT1
Osteoblastic
Chondroblastic
% of genes
60
40
20
0
Gene Lengths
0 - 8077
Short
8077 - 24399
Short Med.
24399 - 66502
Long Med.
> 66502
Long
B
DSRCT
ES
OS
EWS-FLI1 Short Signature
EWS-FLI1 Long Signature
EWS-FLI1 ES Signature
Signature vs. Control Genes (log2FC)
Warm
Cold
Nuclei

**Figure S8. Biases associated gene length in gene signatures. a** The gene set in each sarcoma signature was split into 4 bins of gene length quartiles. **b** EWS-FLI1 ES signature was split by gene length; short < 66502 nt (top row) and long > 66502 nt (middle row). Original gene set without adjustment is in the bottom row.

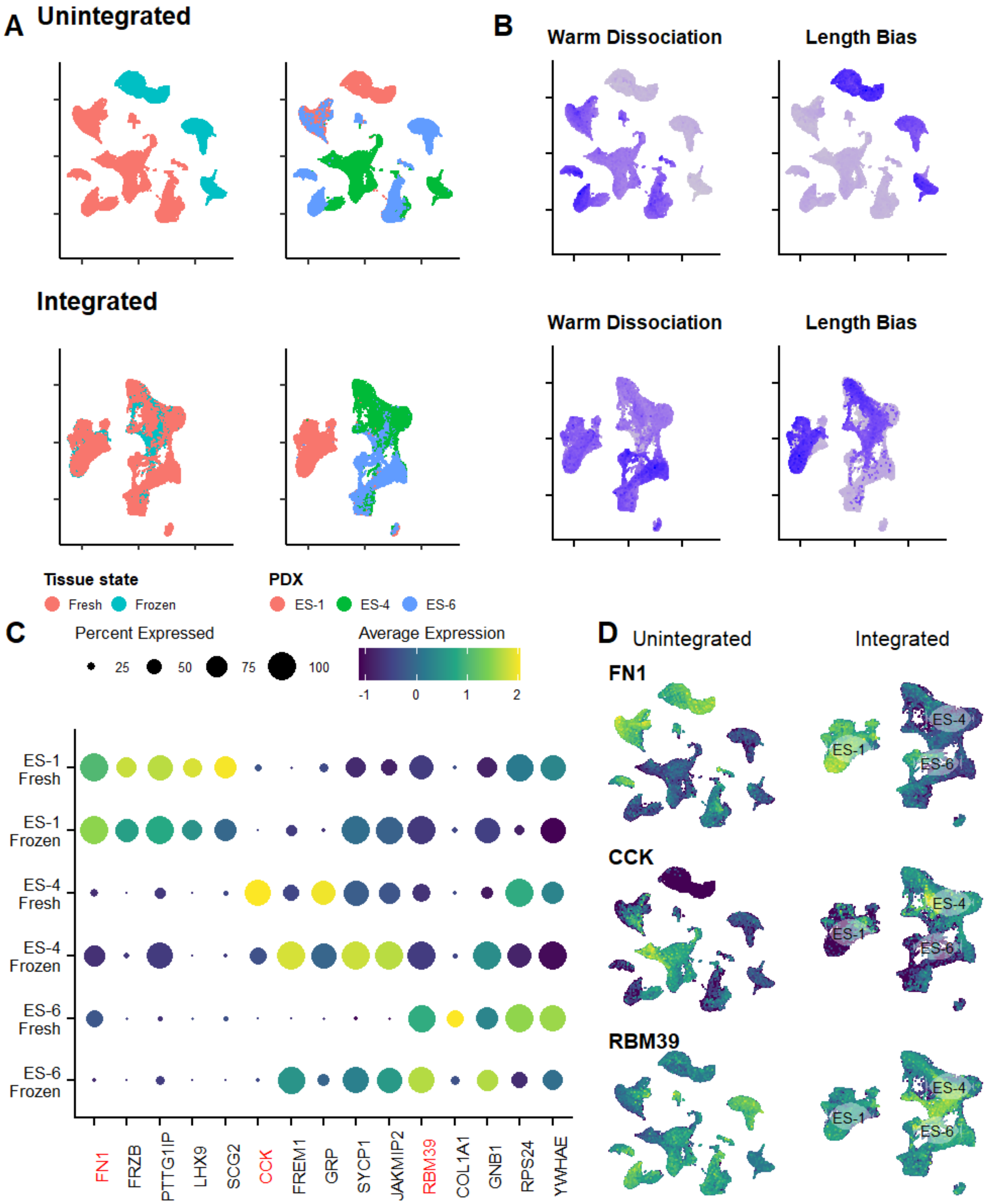


**Figure S9. Integration recovers matching cell states from different dissociation methods for ES. a** UMAP embeddings after integration (bottom row) showed alignment of matching PDX specimens. **b** After integration (bottom row), clusters on the UMAP were no longer affected by the identified biases. Darker blue indicates higher expression. **c** Dot plot of conserved markers show unique gene expression for each PDX. Top marker is in red. **d** After integration (right column), top conserved markers correctly aligned with the associated PDX cluster.

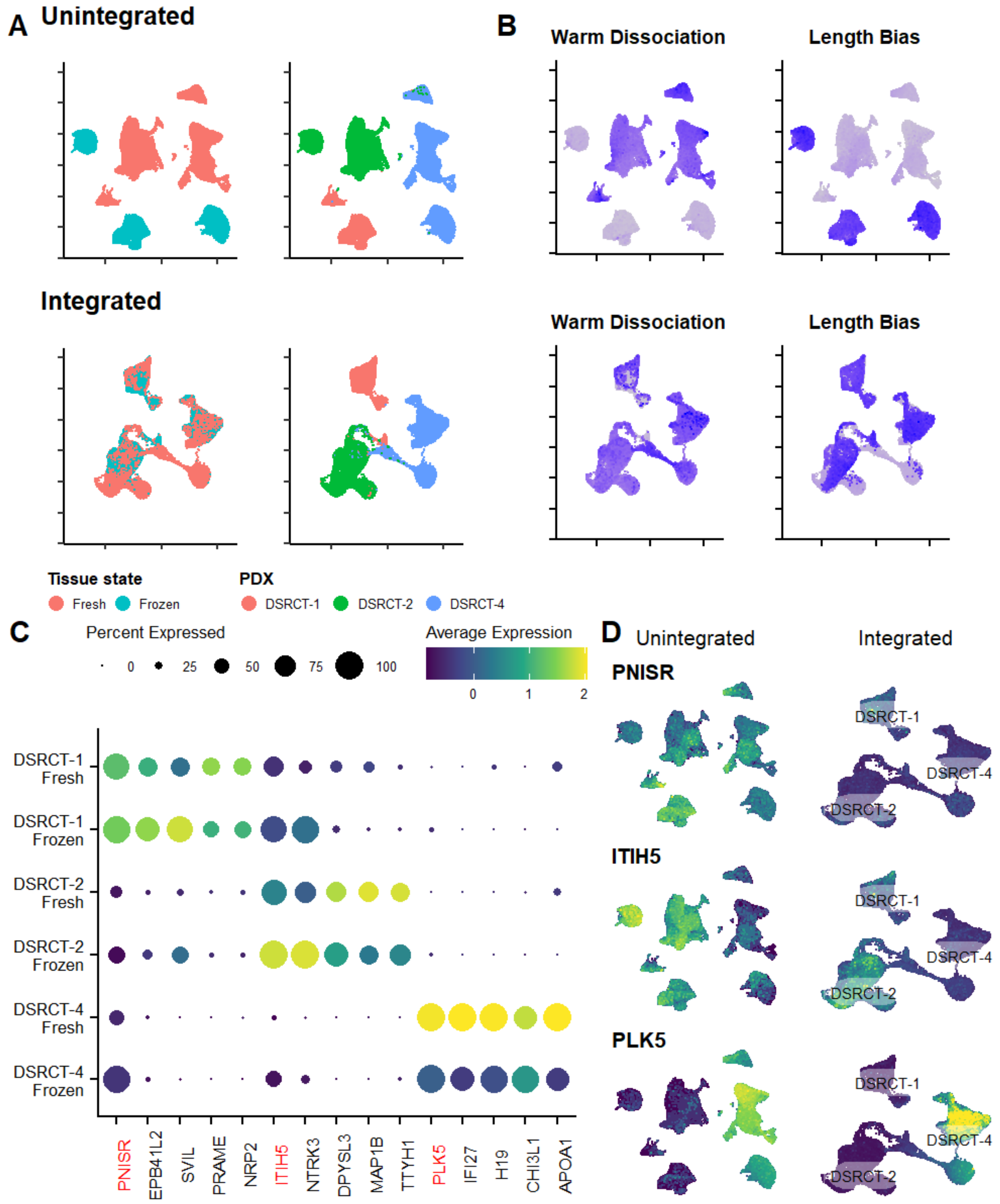


**Figure S10. Integration recovers matching cell states from different dissociation methods for DSRCT. a** UMAP embeddings after integration (bottom row) showed alignment of matching PDX specimens. **b** After integration (bottom row), clusters on the UMAP were no longer affected by the identified biases. Darker blue indicates higher expression. **c** Dot plot of conserved markers show unique gene expression for each PDX. Top marker is in red. **d** After integration (right column), top conserved markers correctly aligned with the associated PDX cluster.

### Additional Tables

**Table S1.** Differentially expressed genes between Warm and Nuclei protocols for ES.

**Table S2.** Differentially expressed genes between Warm and Nuclei protocols for OS.

**Table S3.** Differentially expressed genes between Warm and Nuclei protocols for DSRCT.

**Table S4.** Differentially expressed genes between Warm and Cold protocols for DSRCT.

**Table S5.** Differentially expressed genes between Warm and Cold protocols for ES.

**Table S6.** Top 200 longest genes with the most poly A regions.
